# Unbiased identification of unknown cellular and environmental factors that mediate eQTLs using principal interaction component analysis

**DOI:** 10.1101/2022.07.28.501849

**Authors:** Martijn Vochteloo, Patrick Deelen, Britt Vink, BIOS Consortium, Ellen A. Tsai, Heiko Runz, Sergio Andreu-Sánchez, Jingyuan Fu, Alexandra Zhernakova, Harm-Jan Westra, Lude Franke

## Abstract

Expression quantitative trait loci (eQTL) can reveal the regulatory mechanisms of trait associated variants. eQTLs are highly cell-type and context-specific, but often these contexts are unknown or not measured. Here, we introduce PICALO (Principal Interaction Component Analysis through Likelihood Optimization), an unbiased method to identify known and hidden contexts that influence eQTLs. PICALO uses expectation maximization to identify latent components, referred to as Principal Interaction Components (PIC), that interact with genotypes to maximize explained eQTL effect-sizes.

We applied PICALO to bulk RNA-seq eQTL datasets in blood (n=2,932) and brain (n=2,440). We identify 31 PICs in blood, interacting with 4,169 (32%) unique cis-eQTLs (BH-FDR≤0.05). In brain, we identified 21 PICs, interacting with 4,058 (39%) unique cis-eQTLs (BH-FDR≤0.05). These PICs are associated with RNA quality, cell type composition or environmental influences. Furthermore, PICs clearly disentangle distinct eQTL contexts, for example technical from non-technical factors. Combined, 3,065 unique genes showed a cis-eQTL effect that is dependent on a cell type or other non-technical context, emphasizing the value of methods like PICALO. PICALO is robust, works well with heterogeneous datasets, yields reproducible interaction components, and identifies eQTL interactions and contexts that would have been missed when using cell counts or expression based principal components.

Since PICALO allows for the identification of many context-dependent eQTLs without any prior knowledge of such contexts, this method can help to reveal and quantify the influence of previously unknown environmental factors that play a role in common diseases.

## Introduction

Expression quantitative trait locus (eQTL) mapping is often used to make inferences on the transcriptional consequences of genetic variants that have been identified through genome-wide association studies (GWAS). A challenge of eQTL studies is that the regulatory potential of a variant is often context dependent, resulting in differences in eQTL effects strengths between tissues^1^, cell types^2^, and stimulations^3^. This hinders proper interpretation of disease associated variants^4^. Many different strategies have been employed to identify context dependent eQTLs: for instance, the GTEx consortium generated data in many different tissues^5^ and populations^6^, and in the MetaBrain project many brain eQTL datasets were combined to improve the ability to identify brain dependent eQTLs^7^. Furthermore, single cell RNA-sequencing has been instrumental in identifying cell type dependent eQTLs within the same tissue^8–10^. By using in in vivo stimulations^3, 11, 12^ or case control^13, 14^ comparisons of eQTL effect strength it is also possible to identify stimulation dependent regulatory effects. However, the number of available contexts in these studies has remained somewhat limited, leaving many context-dependent eQTLs to be discovered.

Since data generation is expensive, various computational methods have been developed that do not rely on directly measured contexts or stimulations, but instead estimate factors that influence context dependent eQTLs. The influence of such a context on eQTL effect sizes is often determined through interaction eQTL (ieQTL) analysis^15^. This principle has been applied previously to identify cell type dependent eQTLs, for instance by using predicted cell count measurements in bulk data to estimate the contribution of different cell types to an eQTL effect^15, 16^. More complex models, such as sn-spMF, are capable of detecting factors representing tissue specificity of eQTL effects by using bulk data from different tissues^17^. To identify eQTLs dependent on contexts other than cell types or tissue, other genes have previously also been used in ieQTL analysis. For example, blood-based gene expression levels of other genes were used previously to identify context dependent effects^18^, some of which were related to type 1 interferon signaling. A recent study using the GTEx dataset also revealed context dependent eQTLs that could be attributed to transcript factor levels^19^. The limitations of these methods are that not all confounding contexts might be known or easily measurable, and that individual (gene expression) measurements might not be perfect proxies for specific contexts, and thus can be noisy: for instance, cell type quantifications can differ, depending on the used technology and gate settings whereas measured expression levels of specific genes are unlikely to perfectly reflect environmental stimuli or transcription factor activity. Finally, unbiased methods can also be used to identify ieQTLs for contexts that have not been directly measured or are not readily predicted in a bulk dataset. For instance, by applying principal component analysis (PCA) on gene expression data, it is possible to capture broad transcriptional changes in individual principal components (PCs). Such PCs can then be tested as a potential proxy of a context that might influence eQTL effect strengths. However, PCs often capture gene expression variance explained by a mixture of different biological and technical signals in a single component^18^. Therefore, it is often unclear how to interpret the eQTLs that interact with such PCs.

To attempt to resolve these issues, we developed PICALO (Principal Interaction Component Analysis through Likelihood Optimization), which is an unsupervised method to identify latent components, referred to as Principal Interaction Components (PICs), that serve as proxies for contexts. We applied PICALO to bulk RNA-seq eQTL datasets in blood (n=2,932) and brain (n=2,440). We identified a set of highly informative PICs that together influence >39% of eQTLs. We show that PICs less often capture a mixture of biological and technical contexts, as compared to expression PCs. The observed PICs are associated with RNA quality, cell type composition, and environmental influences, and can be replicated across ethnicities. As such, PICALO is an unsupervised method to identify optimal proxy variables for hidden contexts that influence eQTL effect sizes.

## Results

### PICALO identifies eQTL context PICs through optimization of the interaction log- likelihood, which are less affected by technical confounders than PCs

To identify the context that influences eQTL effect sizes (e.g.: RNA quality, cell type composition, environmental factors), we use the genotype data and tissue expression data for known eQTLs in the relevant tissue (Fig. 1A). Using this data, PICALO identifies the biological and technical context specific to the supplied data by mapping ieQTLs (Fig. 1B) and subsequently optimizing the likelihood of the interaction terms using an expectation maximization (EM) approach (Fig. 1C). In short, an initial guess of a context (such as expression PCs, marker gene expression, cell proportion, etc.) is used to identify an initial set of Benjamini-Hochberg FDR^20^ (BH-FDR≤0.05) significant ieQTLs. If more than one initial guess for a context is supplied (e.g.: multiple different expression PCs), the optimization is started with the initial guess that had the highest number of ieQTLs. Next, the values of this initial guess are adjusted to maximize the log-likelihood of the interaction model for the set of included ieQTLs (this requires at least two ieQTLs). This adjusted vector is subsequently used to reidentify a set of significant ieQTLs (likely larger in size than the set of ieQTLs that were found in the initial ieQTL identification step), after which the process repeats until the vector does no longer change and thus convergence is reached. The resulting context vector is referred to as a PIC (Fig. 1D). The gene expression data is then adjusted for this PIC and its interaction effect with eQTLs using ordinary least squares (OLS)-regression, after which a second PIC can be identified. This procedure is repeated, until no additional significant ieQTLs can be found. The code for PICALO and an animated explanation of the method is available at https://github.com/molgenis/PICALO. A more detailed description of the PICALO method is outlined in the methods section.

**Fig. 1.**
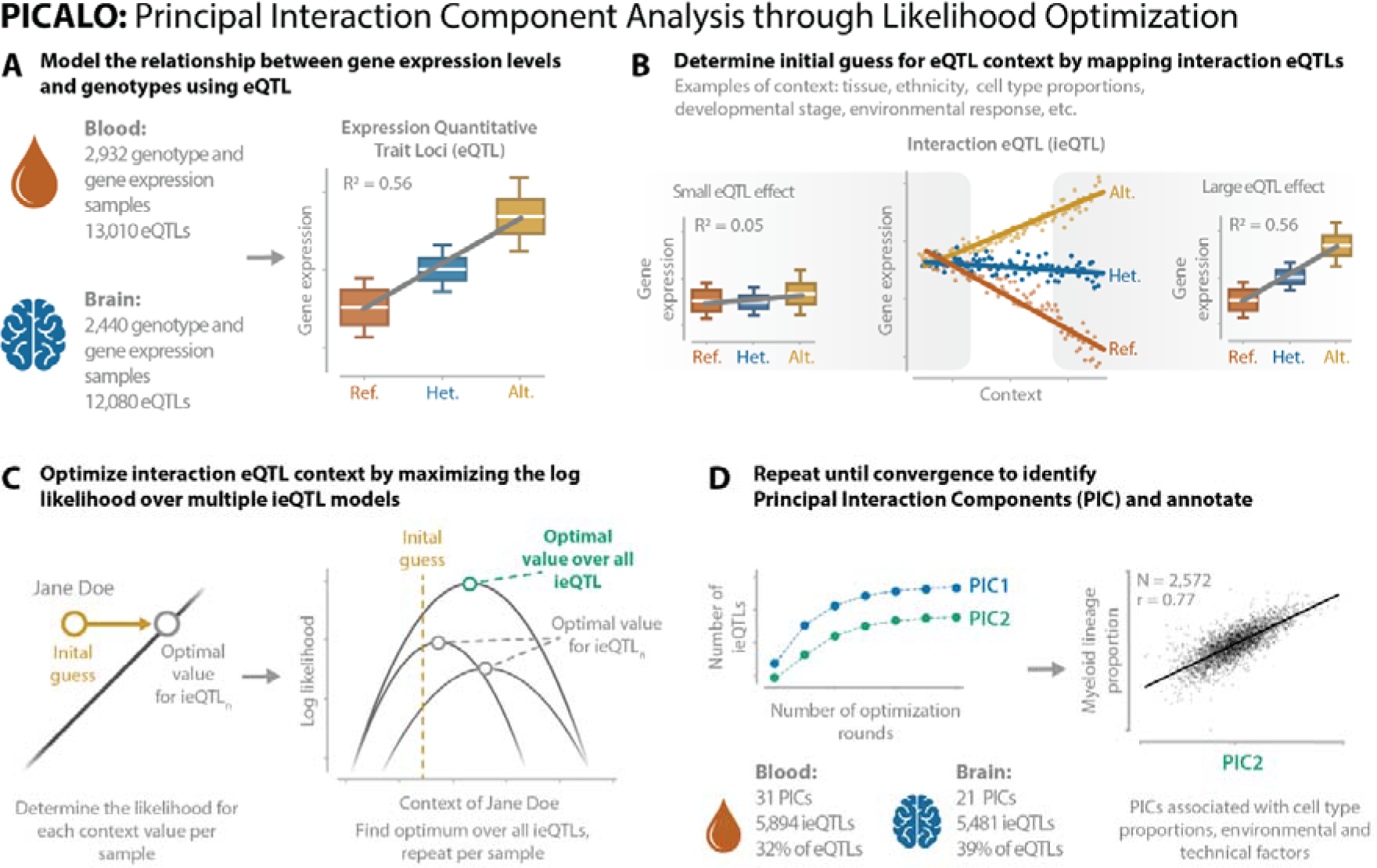
Graphical overview of the PICALO method. (A) PICALO takes eQTL data (i.e.: gene expression and genotype dosage values) as input and (B) maps interactions with a given initial guess of biological / technical context. (C) The initial guess is optimized by maximizing the joint log likelihood on a per sample basis over multiple ieQTLs. (D) Mapping of the interactions and the subsequent optimization are repeated until convergence. The influence of the resulting principal interacting component (PIC) is removed from the gene expression data and the process is repeated until no additional PICs and ieQTLs are identified. The resulting PICs capture technical and biological contexts such as cell type proportions.

We applied PICALO to bulk RNA-seq eQTL datasets in blood (BIOS^18^) and brain (MetaBrain^7^; Fig. 1A). In short, the blood dataset is a large-scale effort from various biobanks in the Netherlands, containing genotype and gene expression data from peripheral blood of population-based samples.

The brain dataset is a large-scale meta-analysis of previously published human brain datasets, including multiple brain regions. In brain we confined ourselves to the cortex samples from European descent. We performed a strict sample selection filter (see methods), to prevent outlier samples, resulting in the inclusion of 2,932 samples in blood and 2,440 samples in brain. We downloaded the summary statistics for the primary eQTLs of each respective study. After filtering, 13,010 eQTLs remained in blood and 12,080 eQTLs in brain.

We first log_2_ transformed the uncentered gene expression data and adjusted the expression data for known technical covariates such as sex, genotype multidimensional scaling (MDS) components, and dataset indicator variables while retaining the mean and standard deviation. Since we observed high correlations between RNA-seq alignment metrics and cell type proportion (Fig. S1), we did not correct for these metrics as this would remove part of the cell type signal. We used the first 25 PCs over this matrix as the initial guess of the eQTL context. After applying PICALO we observed a large increase in the significance of the interaction term for a large proportion of ieQTLs, for instance for TUBB2A and C9orf78 (-log_10_ p-value decrease 71.9 and 45.3) in blood (Fig. 2A) and FAM221A and ADAMTS18 (-log_10_ p-value decrease 41.5 and 49.3) in brain (Fig. 2B). We identified 31 PICs in blood having a total of 5,894 significant interactions (BH-FDR≤0.05) with 4,169 unique eQTLs (32%; Fig. 2A- B; Fig. S2A; Table S1; Table S2). In brain we identified 21 PICs having a total of 5,481 interactions with 4,058 unique eQTLs (39%; Fig. 2A-B; Fig. S2B; Table S1; Table S3). Each PIC showed little to no correlation with other PICs (Pearson r≤0.07 for blood and ≤0.04 for brain; Fig. S3) and the majority of ieQTLs interacted with only one PIC (72% in blood and 75% in brain), suggesting PICs capture unique effects. The first five PICs correlated moderately with the initial guesses (average Pearson r=0.64 ±0.13 for blood and 0.51 ±0.08 for brain; Fig. S4), while subsequent PICs showed lower correlations (average Pearson r=0.20 ±0.15 for blood and 0.17 ±0.1 for brain; Fig. S4).

**Fig. 2.**
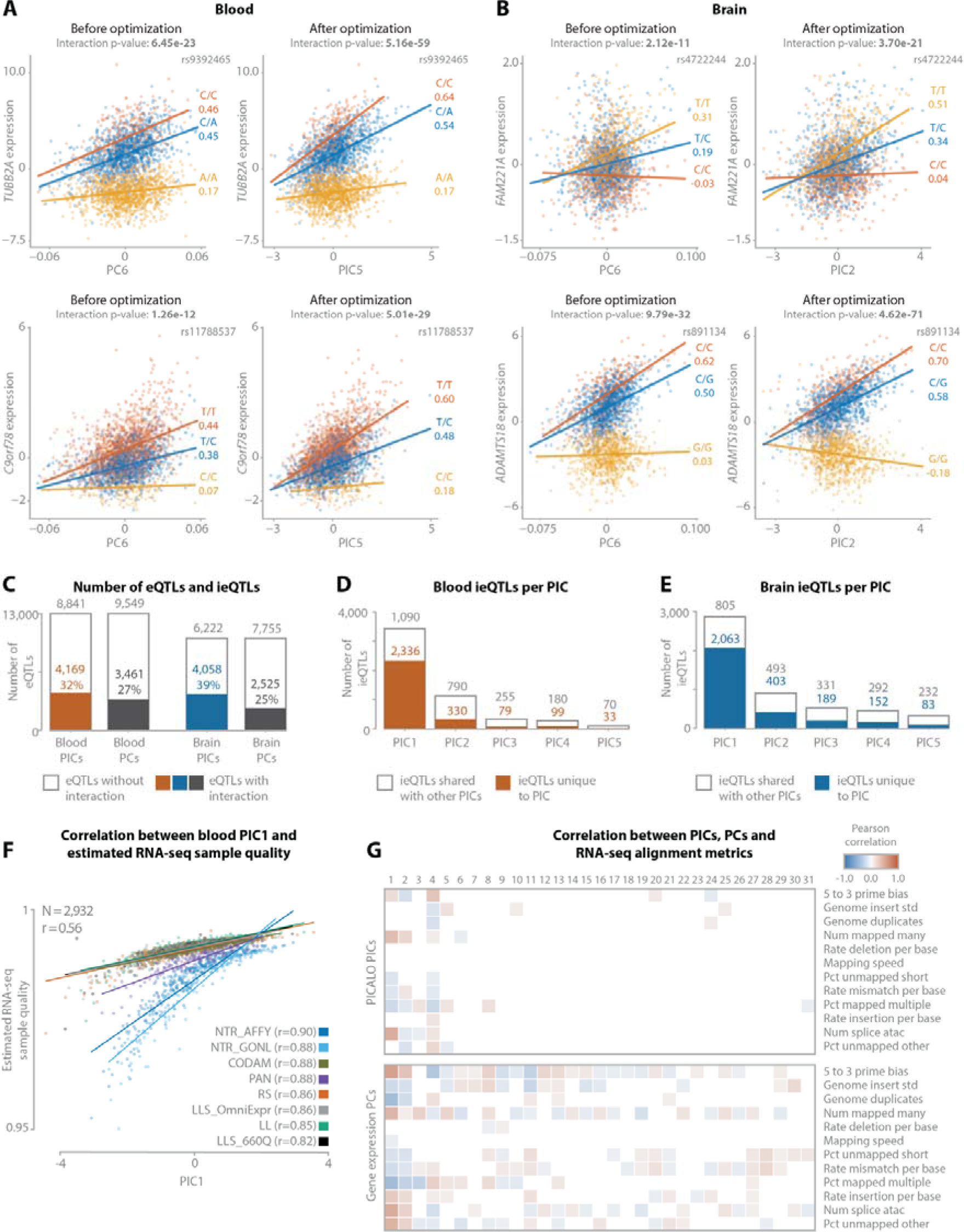
Examples of PICALO optimization for ieQTLs: (A) ieQTL for the genes TUBB2A and C9orf78 with expression PC6 before optimization and resulting PIC5 after optimization in blood, and (B) ieQTL for the genes FAM221A and ADAMTS18 with expression PC6 before optimization and resulting PIC2 after optimization in brain. (C) The number of eQTLs tested in blood and brain and the respective number of eQTLs that have an interaction with one or more PICs or expression PCs. (D) The number of ieQTLs for the first five PICs in blood. (E) The number of ieQTLs for the first five PICs in brain. (F) Regression plot showing the correlation between PIC1 and estimated RNA-seq sample quality calculated as the per sample expression correlation with the overall average expression. (G) Pearson correlation heatmaps correlating PIC (top) and expression PC (bottom) to RNA- seq alignment metrics in blood. The correlation p-values are corrected for multiple testing with Benjamini- Hochberg and only correlations with an FDR≤0.05 are shown. Note that many of the expression PCs correlate significantly with RNA-seq alignment metrics while only a limited number of PICs show significant correlation.

Using an equal number of expression PCs as the number of identified PICs, we found that PICs interact with a higher proportion of unique eQTLs than expression PCs (5.4% increase in blood, 14.9% increase in brain; Fig. 2A). We then evaluated if all variance explained by the interaction terms of the ieQTL models (i.e.: interaction variance) was captured by the identified PICs. For this we adjusted the expression data for the PICs and their interactions and evaluated if we could detect any ieQTLs on the residuals using the first 25 expression PCs (i.e.: the initial guesses on which the PICs were optimized). We observed only a limited number of ieQTLs (7 in blood, 4 in brain), indicating that all interaction variance that is captured by the first 25 PCs is already captured by PICs. We extended this analysis by also evaluating the first 100 PCs, and observed an additional 239 ieQTLs in blood and 40 in brain. This suggests that additional PICs could have been identified if additional expression PCs would have been included as initial guesses, although the number of additional PICs would likely have been limited.

Since EM algorithms can yield results that depend on the choice of the initial starting position (i.e.: initial guess), we evaluated to what extend this was the case for PICALO. We re-optimized the first five PICs in blood using each of the 25 expression PCs as an initial guess independently and compared the results (Fig. S5; Supplementary Note). We observed that PICs capture unique effects that can be robustly identified using PICALO, but that the order in which PICs are identified can be dependent on the initial guess.

### RNA-seq quality and technical confounders are major drivers of eQTL effect sizes in blood and brain

We have shown that PICs are robust proxies for contexts that affect eQTL effect sizes. However, since PICs lack specific annotation, further analysis is required for interpretation. To provide these annotations, we correlated the PICs to known technical factors (e.g.: estimated RNA-seq sample quality, RNA-seq alignment metrics, sample annotations etc.) and biological phenotypes (e.g.: measured or predicted cell type proportions). Subsequently, we evaluated single-cell expression of genes that highly correlate with PICs to further investigate cell type enrichments. Finally, we performed gene set enrichment analysis to investigate pathway and cell type enrichment of positively and negatively correlated genes using the ToppGene suite^21^.

The PIC1 that was identified in blood and the PIC1 identified in brain, individually interact with a substantial number of eQTLs (3,426 in blood and 2,868 in brain). Upon further analysis, both PIC1s showed high correlation with estimated RNA-seq sample quality (blood Pearson r=0.56; Fig. 2F, brain Pearson r=-0.66; Fig. S6A) that we calculated per sample as the correlation between the gene expression per gene and the average expression per gene over all samples. Notably in blood, the correlation per cohort was substantially higher (Pearson r>0.82) than the joint correlation (Pearson r=0.56), suggesting heterogeneity of the estimated RNA quality and consequently that PIC1 provides a more reliable quality measure, perhaps by capturing multiple aspects of technical variation in a single component. Since we did not correct the expression data for RNA-seq quality, it makes sense that PIC1 reflects this.

We then determined how many PICs were affected by RNA-seq alignment metrics and compared this to expression PCs. We calculated the Pearson correlations between each PIC and RNA-seq alignment metrics (e.g. those from PICARD, FastQC; Fig. 2G; Fig. S6B) and observed that 10 out of 31 PICs had a significant PIC-metric correlation in blood (9.4% of pairs; BH-FDR≤0.05) and 7 out of 21 PICs in brain (16.1% of pairs). In contrast, for an equal number of expression PCs we observed that 29 out of 31 expression PCs had a significant correlation in blood (41.7% of pairs) and 21 out of 21 expression PCs in brain (49% of pairs). This indicates that PICs distinguish technical from non-technical factors better than PCs and that most PICs do not reflect technical confounders.

We next classified PICs as being predominantly technical if they were correlated significantly with an RNA-seq alignment metric, but not with one of the cell-count proportions or other biological factors. Using this principle, we determined that PIC1, PIC4, and PIC8 are predominantly technical in blood (Fig. 2G, Fig. 3A) and PIC1, PIC4, and PIC7 in brain (Fig. S6B, Fig. 4A). In total, the technical PICs accounted for 3,760 interactions (64%) with 3,555 unique eQTLs (27%) in blood and 3,416 interactions (62%) with 3,140 unique eQTLs (31%) in brain. The remaining non-technical PICs accounted for 2,134 interactions (36%) with 1,617 unique eQTLs (12%) in blood and 2,065 interactions (38%) with 1,671 unique eQTLs (16%) in brain. In blood and brain combined, we identified non-technical interaction effects for 3,065 unique genes.

**Fig. 3.**
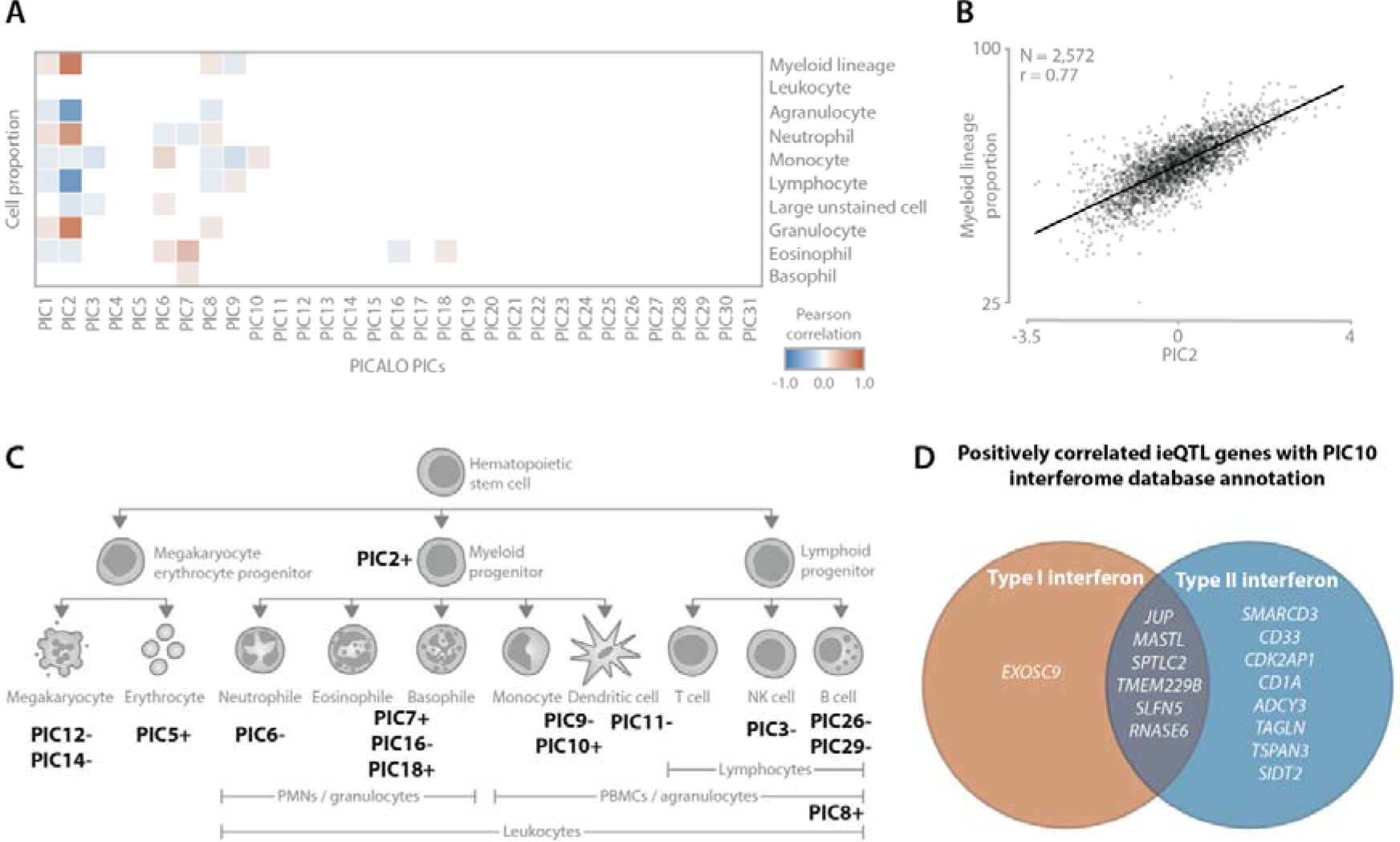
(A) Pearson correlation heatmaps correlating PICs to measured cell type proportions in blood. The correlation p-values are corrected for multiple testing with Benjamini-Hochberg and only correlations with an FDR≤0.05 are shown. (B) Regression plot showing the correlation between PIC2 and myeloid lineage cell proportions (granulocyte + monocyte) in blood. (C) Simplified overview of the blood cell type lineage with annotations of PICs describing distinct (groups of) cell types. Positive and negative signs indicate the direction of effect. (D) Negatively correlating eQTL genes interacting with PIC10 showed enrichment for type II interferon signaling as annotated by the Interferome Database Annotation.

**Fig. 4.**
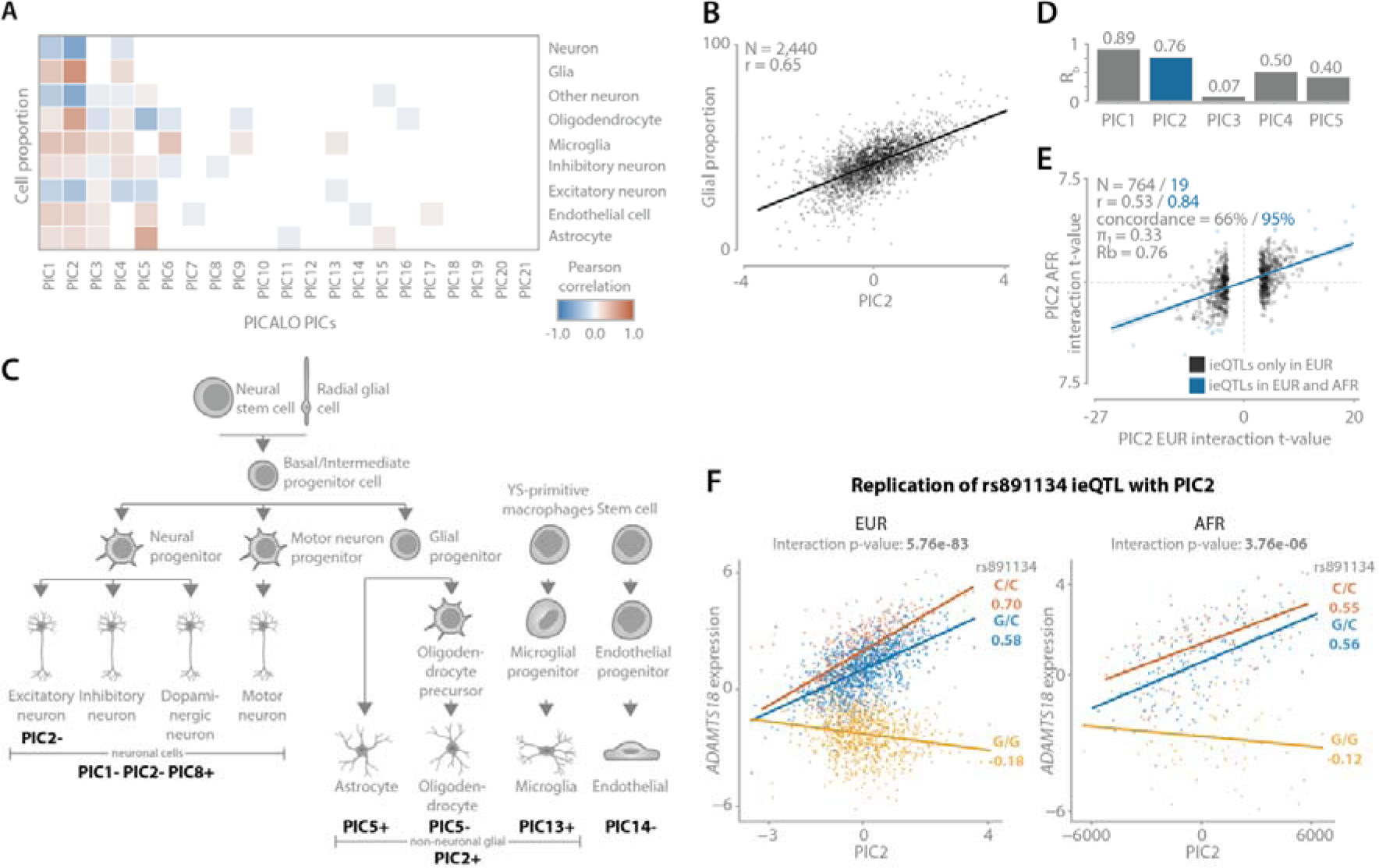
(A) Pearson correlation heatmaps correlating PICs to predicted cell type proportions in brain. The correlation p-values are corrected for multiple testing with Benjamini-Hochberg and only correlations with an FDR≤0.05 are shown. (B) Regression plot showing the correlation between PIC2 and glial proportion (microglia + oligodendrocyte + astrocyte) in brain. (C) Simplified overview of the brain cell type lineage with annotations of PICs describing distinct (groups of) cell types. Positive and negative signs indicate the direction of effect. (D) Rb replication statistics for the replication of eQTLs interacting with the first five PICs discovered in samples of European descent (EUR) and replicated in samples of African descent (AFR). (E) Regression plot showing the interaction t-values of PIC2 ieQTLs discovered in EUR and replicated in AFR. Blue points are significant in both datasets, the statistics of which are shown in blue. (F) Example of replicating ieQTL: rs891134 affecting ADAMTS18 gene expression and interacting with PIC2. The left plot shows the interaction in EUR, the right plot shows AFR. The x-axis shows the PIC2 scores, the y-axis shows the covariate corrected gene expression, each dot represents a sample. The p-values are calculated using the unconditional ieQTL analysis. Colors indicate SNP genotype. Values under the alleles are Pearson correlation coefficients.

### Specific types of cells, interferon signaling, and prior cytomegalovirus infection influence eQTL effect size in blood

To interpret the non-technical PICs in blood, we first correlated their values with measured cell counts and combinations of cell types from the same lineage (Fig 3A, Fig. S7A). For 10 out of 31 PICs we observed a significant correlation (PIC1-PIC3, PIC6-PIC10, PIC16, and PIC18; BH-FDR≤0.05). For example, PIC2, interacting with 1,120 eQTLs, showed the strongest correlation with myeloid lineage cell percentage (Pearson r=0.77; Fig. 3B), consisting predominantly of neutrophils. Furthermore, for PIC7, interacting with 60 eQTLs, we observed a correlation with measured eosinophil proportion (Pearson r=0.36).

We then evaluated whether PICs were enriched for certain cell types and biological processes by correlating the PICs with gene expression levels and splitting the genes into groups of positively and negatively correlating ones. Using the top 200 positively and 200 negatively correlated genes per PIC (Table S4), we tested for gene set enrichments (Table S5) and evaluated expression in purified and single-cell datasets (Fig. S8 and Fig. S9). This analysis further supported the neutrophil association for PIC2 and the eosinophil association for PIC7. Furthermore, we were able to assign putative labels of (sub)types of cells to nine more PICs (Fig. 3C). For PIC5 (103 ieQTLs), for example, we found that the positively correlating genes showed a strong enrichment of erythrocyte-specific genes (ToppCell enrichment p-value <1.4×10^-233^). Moreover, this PIC was enriched for red blood cell pathways, such as for the uptake of oxygen and release of carbon dioxide (Pathway enrichment p-value=6.7×10^-11^), among others (Table S5). We also found that PICs interacting with a small number of eQTLs also showed high specificity for specific cell types. For instance, the negatively correlating genes for PIC26 (6 ieQTLs) and PIC29 (11 ieQTLs) both showed strong enrichment for B-cells (ToppCell enrichment p-value <8.4×10^-173^ for PIC26 and p-value <6.2×10^-147^ for PIC29). Moreover, single-cell expression showed that these genes are specifically expressed in B-cells, further confirming that these PICs capture B-cell differences.

Next, we evaluated if the top 200 positively and negatively correlated genes per PIC were enriched for certain pathways. In the Zhernakova et al. study^18^, which uses the same data as this study, an interaction module was identified that was a proxy for type-1 interferon response. Here, we found that PIC10 was enriched for the interferon signaling pathway (Pathway enrichment p-value=1.6×10^-36^). While the number of affected eQTLs by PIC10 was lower than the number of eQTLs affected by the equivalent module in Zhernakova et al. (44 as compared to 145), the enrichment p-value for the interferon signaling pathway was substantially more significant (p-value=1.6×10^-36^ versus p- value=2×10^-6^). Finally, by using the interferome database annotation^22^, we found that the 21 out of 22 genes that positively correlate with PIC10 were involved in type 2 interferon signaling (Fig. 3D).

Finally, we found a significant correlation (Pearson r=-0.29, p-value=5.97×10^-15^; Fig. S10A-B) between PIC3 and the presence of anti-bodies against cytomegalovirus (CMV). This was determined by a CMV infection signal that was calculated from an aggregate of multiple IgG antibody profiles enriched in a PhIP-seq experiment for 1,433 samples^23^. These CMV anti-bodies can be indicative of a past or latent infection with CMV, which is expected for 45% of the Dutch population^24^. Latent CMV infection can have a lasting effect on the immune system^25^, increases monocyte differentiation^26^, and CMV can reprogram monocyte gene expression^27, 28^. We also observed a correlation between the CMV antibodies and expression PC5 (r=-0.29, p-value=1.14×10^-15^). Although PIC3 and PC5 seem very similar on first glance (Fig. S10C; Pearson r=0.66), we observed that PC5 was correlated with multiple cell type proportions (max absolute Pearson r=0.27 with neutrophil proportion; Fig. S10D), while PIC3 was not (max absolute Pearson r=0.14 with monocyte proportion; Fig. S10D). While this suggests that the CMV infection context can also be captured by using expression PCs, it also suggests that eQTL interactions with such a context would be biased by cell type composition differences, which is not the case for the detected PIC.

### Specific types of cells influence eQTL effect size in brain

In contrast to the PICs in blood, the PICs identified in brain were harder to annotate since no measured cell type proportion measurements were available for this dataset. However, by using the predicted cell type proportions as used in de Klein et al. (Fig. S7B), and gene set or pathway enrichment analysis (Table S5) as well as a comparison with brain single-cell expression from ROSMAP (Fig. S9B), we were able to assign putative labels to a total of nine PICs.

First, PIC2, which has a significant interaction with 896 eQTLs, correlated strongly with predicted glia proportion (Pearson r=0.65). Second, for PIC5 we identified 315 ieQTLs and strong correlations with predicted astrocyte (Pearman r=0.48) and oligodendrocyte proportion (Pearman r=-0.45) as well as a high enrichment of cell type specific genes for these cell types (p-value <1.5×10^-136^ for astrocyte and p-value <6.2×10^-99^ for oligodendrocyte). Less frequent cell types were also captured by PICs that interact with only a limited set of eQTLs. For PIC13, interacting with 13 eQTLs, while we observed a low but significant correlation with predicted microglia proportion (Pearson r=0.08), we did observe a clear enrichment for microglia in the single-cell data, as well as an enrichment of microglial genes (ToppCell enrichment p-value <1.4×10^-165^). Finally, genes negatively correlating with PIC14, which interacts with 17 eQTLs, showed enrichment for endothelial cells in single-cell data as well as when using gene set enrichment (ToppCell enrichment p-value <1.2×10^-51^). However, similar as PIC13, only a weak correlation was observed between PIC14 and the predicted endothelial cell counts (Pearson r=-0.07).

### Top brain PICs replicate well in an independent dataset

Lastly, we evaluated how well PIC ieQTLs replicate in an independent dataset and calculated the ieQTL agreement using three metrics (i.e.: allelic concordance, π_1_ ^29^, and R ^30^). For this we used the PICs identified in the 2,440 brain samples from European descent and replicated them in 311 brain samples from African descent. We observed that the top two PICs replicate very well (PIC1: allelic concordance=76%, π_1_=0.3, R_b_=0.89; PIC2: allelic concordance=66%, π_1_=0.33, R_b_ = 0.76; Fig. 4D-E; Fig. S11; Error! Reference source not found.). The high concordance of technical PIC1 is most likely due to the overlap of shared confounding effects between samples of European and African descent originating from the same cohorts. As expected, the remaining PICs replicate less well as the number of ieQTLs decreases with subsequent PICs and they capture a lower proportion of interaction variance (Error! Reference source not found.). For the biological PIC2, annotated as glial cell proportion, a total of 10 ieQTLs replicated. This included, ADAMTS18 (Fig. 4F), which has previously been identified as oligodendrocyte specific^31^. This finding supports our earlier observation that PIC2 captures glial proportion differences.

## Discussion

We have developed PICALO, a novel method that uses an EM algorithm to identify unique principal interaction components that affect eQTL effect sizes. We identify 31 PICs in brain, and 21 PICs in blood that together significantly interact with up to 39% of the included eQTLs, reflecting thousands of genes, and which is a substantial improvement over Zhernakova et al. who reported significant interactions for only 12% of the eQTLs^18^. While we could also assign a biological label to thousands of ieQTLs, many of the identified eQTLs interacted with technical PICs. This could suggest that for a large proportion of eQTLs the effect size depends on technical covariates even after stringent correction. We therefore reason that the technical PICs identified by PICALO can be used to remove additional technical variance from eQTL studies to further improve eQTL effect size estimates, which in turn might improve downstream analyses with GWAS signals such as colocalization or Mendelian Randomization. Overall, the identified PICs describe highly relevant biological and technical eQTL contexts without knowing them a priori. Moreover, they provide better differentiation between technical and biological components as compared to PCs.

Application of PICALO is not limited to bulk RNA-seq and could also be applied in single-cell data to identify stimulation contexts and specific types of cells including the less frequent ones. We have shown that expression PCs can be used as an initial guess of eQTL context, but we do note that it is also possible to start with marker gene expression levels or cell-type proportions. We expect that PICALO can also be applied to other quantitative phenotypes, such as clinical data, ethnicity information, or other molecular phenotypes such as protein levels. While we have focused on cis-eQTLs in this study, PICALO could also be applicable to trans-eQTLs. For many trans-eQTLs it is currently unclear whether they are the consequence of cell type proportion differences or actual regulatory effects^32^, a distinction that can potentially be improved using a combination of PICs identified by PICALO.

A limitation of PICALO is that the identified PICs do not have a direct functional annotation and therefore require further analyses for proper interpretation, which we have shown to be feasible. Furthermore, we have shown that PICs are uncorrelated to each other and capture unique effects influencing eQTL effect sizes. However, like expression PCs, PICs may capture shared effects between distinct biological or technical processes within a single PIC, possibly complicating the interpretation. Another limitation is that lowly expressed genes cannot be included as their interactions cannot be properly corrected for. Finally, we note that PICs are currently restricted to linear effects with genotype.

To sum up, PICALO is useful for the analysis of eQTL datasets to detect the relevant contexts that influence eQTLs. Currently our method is especially well-powered in large sample sizes, but various large-scale eQTL and pQTL studies are currently underway or have just been completed^32, 33^. Therefore, we believe PICALO will prove highly useful in the future, enabling the discovery of previously unknown contexts that influence eQTL effect sizes, and consequently potentially improving the interpretation of disease associated variants.

## Methods

### General description of PICALO

PICALO is an EM based algorithm for the unbiased identification of known and unknown contexts that influence the regulatory effect of genetic variants on gene expression. PICALO takes as input a set of eQTLs and their corresponding expression and genotype data, as well as a set of initial guesses (e.g.: potential context components). Optional arguments allow for the correction of technical covariates both with and without an interaction term. PICALO can deal with missing genotypes and can work with multiple, heterogeneous eQTL datasets. In this paper we show the use of expression PCs as a starting point for potential contexts, however, other quantitative phenotypes or characteristics such as cell type fractions or marker genes can also be used. Using this data, PICALO identifies latent components that maximally affect eQTL effect-sizes (i.e.: PICs) in three steps. These steps are detailed below.

### Step 1. eQTL inclusion criteria and data pre-processing

PICALO allows for the filtering of eQTLs based on the following metrics: eQTL p-value (default ≤0.05), genotype call rate (default >0.95), number of samples per allele (default >2), minor allele frequency (MAF; default >0.05), and Hardy-Weinberg equilibrium p-value (default >1×10^-4^). If the input data consists of multiple datasets, the call rate is calculated per dataset and all samples of the same dataset are considered missing if the call rate is not met. Note that eQTLs for which the included samples have a low average expression (e.g.: log_2_(TMM +1) ≤1) should be manually removed prior to applying PICALO.

### Step 2. Correcting the gene expression levels for technical confounders

For the gene expression, PICALO expects the input to be log_2_-transformed, centered per gene, and z-transformed per sample over all genes. To correct for technical confounders, PICALO constructs a design matrix encompassing all supplied technical covariates and automatically includes dataset indicator variables if more than one dataset is used. An extra term for the interaction between a technical covariate and genotype can be included as well. If applicable, dataset indicator variables always include a term for the interaction with genotype to correct for possible dataset specific interactions. Before the identification of each PIC, the input gene expression data is corrected for all terms in the design matrix using OLS-regression in which samples with missing genotypes are ignored. The residuals are used as gene expression input for step 3.1. After identification, each PIC is included in the design matrix with a term for the interaction with genotype.

### Step 3.1 Expectation maximization: identification of interaction eQTLs

As the first part of the EM step, PICALO identifies the starting point that has the highest number of significant ieQTLs. A graphical overview of this step is shown in Fig. S12A. The starting point is an initial guess of the eQTL context and is hereafter referred to as context. The significance of an ieQTL is analyzed as follows: first the genotype and context effects are removed from the expression data, ensuring that only the interaction term explains any variance in the model. PICALO then forces the distribution of the gene expression levels and the context(s) into a normal distribution per dataset by ranking with ties. Per eQTL, the significance of the interaction term is calculated by comparing the residual sum of squares (RSS) of two linear models: one without (eq. 1) and one with (eq. 2) the interaction term included.

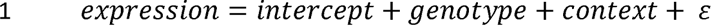

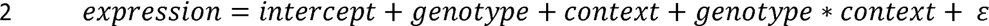

The interaction p-value is calculated using a one-sided F-statistic with (1, n-4) degrees of freedom after which BH multiple testing correction is applied per context. Only ieQTLs that are significant (BH-FDR≤0.05) are used for the optimization step. If there are less than two ieQTLs available for optimization for any of the samples, the program is halted.

### Step 3.2 Expectation maximization: optimization of interaction component

For the optimization of the context, PICALO maximizes the log-likelihood of the interaction model (eq. 3) over all significant ieQTLs considering each sample individually. A graphical overview of this step is shown in Fig. S12B. For clarity, first consider the case in which only one ieQTL is optimized. Using the beta estimates as calculated in eq. 2, the log-likelihood of these beta values, given the observations (e.g.: gene expression levels, genotype, context) can be calculated as follows:

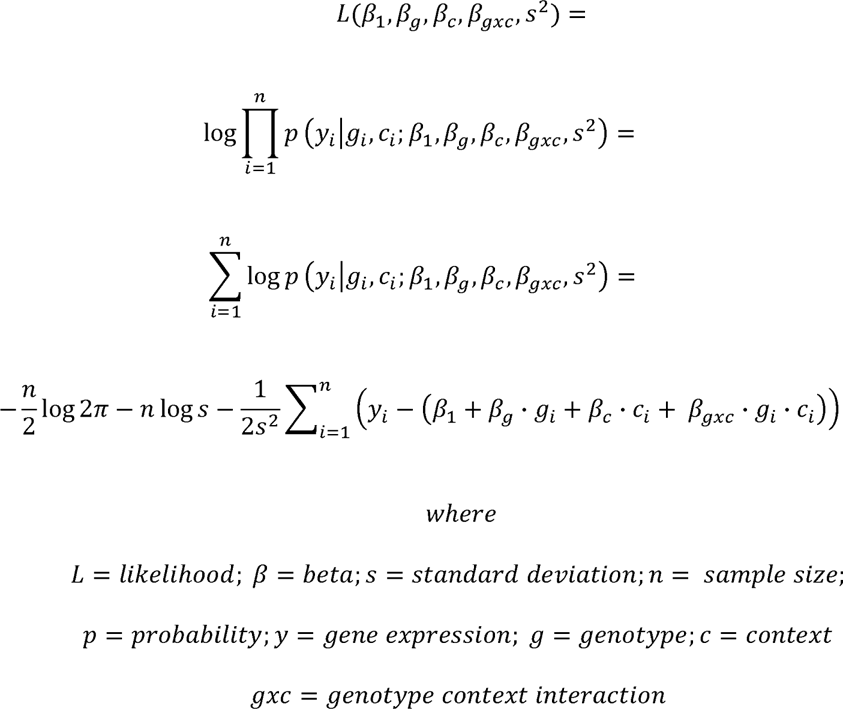

In other words, the likelihood of the model equals the joint probability of the model residuals coming from a normal distribution with mean zero and a standard deviation of s^2^. In contrast to how the likelihood is traditionally applied (i.e.: likelihood of model parameters given the data), PICALO calculates the likelihood of the context value for a single sample given the model parameters. The underlying assumption is that the context estimate of a given sample has a certain margin of error but that this error has an average of zero over all samples (i.e. the model parameters are unbiased). PICALO calculates the log-likelihood of sample_i_ having context value c_i_ given that µ_c_ and µ_gxc_ are true. In other words, PICALO adjusts the value for c_i_ to maximize the log-likelihood of the complete model while all other observations and parameters, especially all values for c” c_i_, are kept fixed. In the case of a single ieQTL, this translates to adjusting the context value of each sample (Fig. S13A) and determining the change in log-likelihood (Fig. S13B). In principle, the log-likelihood is maximal for a given sample when its context value intersects with the regression line of its corresponding genotype group.

We assume s^2^ to be constant since we forced the expression and context distributions into a normal distribution per dataset by ranking with ties (see step 3.1). As a result, maximizing the log-likelihood equals to minimizing the RSS of the model (eq. 4).

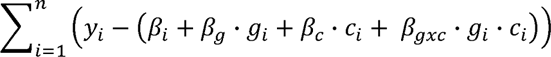

Extending this concept to m ieQTL models, we can find the joint maximum log-likelihood by identifying the context value that gives the lowest RSS over all ieQTLs. This optimum can be calculated efficiently by describing the log-likelihood per sample and per model as coefficients of a second-degree polynomial, summing these together per sample, and finally identifying the focus of the joint function (Fig. S13B).

We then repeat this process by using the optimized context as input context at the start of step 3.1. Note that in the second round the optimized context is always used as starting context for optimization, therefore, the initial guess selection step is skipped. If the minimum number of iterations has been reached (default 5), and the Pearson correlation between the current and previous optimized context vectors is above the tolerance (default ≥0.999), the covariate is considered converged and the loop is terminated. This converged covariate is referred to as a PIC. In some cases, the optimization step can get stuck in an oscillating loop in which the values of the context in subsequent iterations are reverted to those of previous ones. If this occurs, the context is also considered converged and the iteration (e.g. current or previous) with the highest number of ieQTLs is returned as PIC. Step 2 and 3 are repeated until the required number of PICs are identified or <2 ieQTLs are available for optimization. Note that an initial guess can be used more than once to derive a PIC. As a result, the number of PICs that PICALO identifies are not limited to the number of initial guesses that are supplied.

### Data pre-processing

We made use of the bulk RNA-seq eQTL datasets collected by BIOS^18^ (peripheral blood; n = 3,997) and MetaBrain^7^ (multiple brain regions; n=8,727). For BIOS, we confined ourselves to the samples included in eQTLgen (n-samples=3,831, n-cohorts=9)^32^, and used the RNA-seq data as aligned and pre-processed for that project. For MetaBrain, we confined ourselves to the cortex samples of European descent (EUR; n-samples=2,683; n-cohorts=14) and African descent (AFR; n-samples=319, n-cohorts=3). The EUR samples were used for discovery and the AFR samples for replication and were processed independently. In short, all samples were realigned in the same manner and the genotyping array data was imputed using the Haplotype Reference Consortium (HRC) panel. Details on the quality control can be found in the respective manuscripts.

### Genotype data processing

We first ran MixupMapper^34^ to identify sample mix-ups in BIOS. In short, MixupMapper identifies cis-eQTLs on a dataset and determines if samples are more often an outlier than could be expected by chance. This resulted in 15 possible sample mix-ups that were excluded from the BIOS dataset. For MetaBrain EUR we removed 223 genetically similar individuals (pihat >0.125) as calculated with PLINK 2.0^35^. Furthermore, we excluded all samples for which RNA-seq alignment metrics or sex information was unavailable, as well as datasets with less than 30 samples. Over the remaining samples we determined the first four MDS components over the genotype data using PLINK. We included variants that passed the following thresholds: MAF >0.1, Hardy-Weinberg exact test p-value >0.001, missingness per variant ≤5%. We then pruned the SNPs using the independent pairwise setting with windows size of 1000 kb, step size of 50 and R^2^ threshold of 0.5. In total 289,059 variants for BIOS and 52,635 variants for MetaBrain EUR passed pruning and were used to identify the first four MDS components (Fig. S14). In BIOS we then excluded a total of 20 samples that had a z-score >3 standard deviations on one of the four components. For MetaBrain EUR we observed that ENA samples formed a separate cluster (Fig. S15), most likely because these samples have RNA-seq derived genotypes. While these genotypes are reliable and can be used for local imputation and eQTL mapping, we found that the non-genome wide coverage induced batch effects when applying PICALO. We therefore excluded this dataset consisting of 243 individuals. In the MetaBrain AFR samples we excluded 8 outlier samples. Finally, 2,932 samples remained in BIOS, 2,440 samples in MetaBrain EUR, and 311 samples in MetaBrain AFR for which we reidentified the genotype MDS components to use as technical confounder (Fig. S16A and B).

### Initial eQTL context prediction and gene expression pre-processing

We used the Trimmed mean of M values (TMM) normalized expression data for BIOS and MetaBrain separately. First, we estimated an initial guess of the eQTL context required by PICALO by performing PCA over the uncentered, but covariate corrected gene expression matrix. This procedure was as follows: we selected the included samples, removed genes with no variation, log_2_- transformed, and saved the mean and standard deviation per gene. As covariates we used dataset indicator variables, sex, and the four genotype MDS components which we removed using OLS-regression. We then restored the mean and standard deviation in the residuals, as calculated per gene prior to the covariate correction and then performed principal component analysis (PCA) analysis (Fig. S16C and D). The first 25 expression PCs were saved as initial guesses for PICALO.

We then pre-processed the gene expression matrix to use as input for PICALO. For this, we took the TMM expression data for BIOS and MetaBrain (performed separately) and applied the following steps: we selected the included samples, removed genes with no variation, performed log_2_- transformation, centered and scaled the genes and z-score transformed the samples.

Last, we prepared the gene expression matrix to be used for gene set enrichment analysis. For this we took the gene expression matrix as prepared above (i.e. TMM log_2_-transformed, centered and scaled, z-transformed) and removed the included technical covariates (i.e. , sex, four genotype MDS components, and dataset indicator variables) from the expression data using OLS-regression. To verify that there was no major residual between-datasets variance we performed PCA and observed that datasets clustered together (Fig. S16E and F).

### eQTL data processing

We next downloaded eQTLs for BIOS and MetaBrain from <u>molgenis26.gcc.rug.nl</u> and <u>metabrain.nl</u>. We only included primary eQTLs and excluded meta-gene eQTLs. We further removed eQTLs for which the genes had an average expression (TMM log_2_+1) <1 in our sample subset, or for which the SNP had a MAF ≤5%, Hardy-Weinberg exact test p-value <0.0001, or those that had less than two samples in each genotype group. This left 13,010 eQTLs for BIOS and 10,280 eQTLs for MetaBrain that we used for downstream analyses.

### PICALO analysis

We determined the optimum number of expression PCs to correct for by testing 0 up to and including 100 expression PCs in steps of 5 and determined the number of expression PCs that maximizes the number of found ieQTLs. This resulted in 40 expression PCs for blood and 80 for brain that were included as technical covariates (note that the first 25 PCs of these sets were used as initial guess of the eQTL context). We further included dataset indicator variables, sex, and the first four genotype MDS components including their interactions with genotype as technical covariates. We then ran PICALO with a minimum of 50 iterations and a maximum of 100 iterations. We found that forcing a higher minimum number of iterations (default is 5) can in rare situations result into more robust PICs as the program has more time to climb out of local minima. We do note that in the majority of the cases, the model converged within 20 iterations.

### Detection of eQTL interacting with PICs / expression PCs

In order to make a fair comparison between the number of ieQTLs with PICs versus expression PCs, we remapped these interactions separately from PICALO similar as outlined in the methods section step 1 and step 3.1. However, in this case, we did not remove the genotype and covariate effects from the expression data prior to performing the F-test. Moreover, we performed this as a conditional analysis: first we determined the ieQTLs significant for the first input context (PIC or expression PC). Then, we removed the first input context and its interaction with genotype from the gene expression levels using OLS-regression. Finally, we repeated this process for subsequent input contexts, where all previous input contexts were also included in the OLS-regression correction. This conditional analysis ensures that the number of ieQTLs is not inflated due to correlated covariates.

### Gene set enrichment analysis

In order to annotate the PICs we performed functional enrichment of cell types and pathways using the ToppGene functional analysis tool^21^. For this we used the covariate corrected gene expression matrix and forced the expression distribution per gene into a normal distribution per dataset by ranking with ties. We removed genes that had an average expression (TMM log_2_+1) <1 in our sample subset and calculated the Pearson correlation between each PIC-gene pair. Two sets of gene enrichment analyses were performed: (1) over the top 200 genes with the highest or lowest significant (BH-FDR≤0.05) Pearson correlation z-score, and (2) over the ieQTL genes. In both cases the results were split in positively and negatively correlating genes. Standard ToppGene settings were used, and only significant results were reported (FDR≤0.05).

### BLUEPRINT purified cell expression data

We downloaded the BLUEPRINT^36^ bulk RNA-seq expression of purified venous blood and cord blood fractions from http://dcc.blueprint-epigenome.eu. In case of multiple experiments on the same cell type we took the mean Fragments Per Kilobase of transcript per Million (FPKM) value per gene. Genes with a mean FPKM smaller than one and that showed no expression in at least 90% of the cell types where excluded. We log_2_ transformed the FPKM+1 values and performed a center and scaling per sample.

### ROSMAP brain single cell expression data

We downloaded the ROSMAP^37^ single-nucleus RNA-seq data from Synapse. This dataset encompassing 80,660 single-nucleus transcriptomes from the prefrontal cortex of 48 individuals with varying degrees of Alzheimer’s disease pathology. The pre-processing of the data is described in de Klein et al. 2021. In short, we normalized the expression matrix on a per individual per cell type basis. We then created expression matrices for each broad cell type (excitatory neurons, oligodendrocytes, inhibitory neurons, astrocytes, oligodendrocyte precursor cells, microglia, pericytes, and endothelial cells) by calculating the average expression per gene and per individual. We included only genes that showed expression in at least 90% of the cell types and then performed a center and scaling per sample.

### Cell type expression enrichment of PICs

The BLUEPRINT and ROSMAP expression in different blood and brain cell types was used to show that the genes associated to PICs show cell type specific expression patterns. The same top 200 associated genes top a PIC were used as for the gene set enrichment analysis. Per cell type we calculated the mean expression of the associated genes to further show how the PICs reflect cell type composition differences.

### Replication of brain PIC ieQTLs in an independent dataset

In order to test if PIC ieQTLs are robust across datasets, we replicated the ieQTLs identified in brain (MetaBrain EUR; n=2,440) in the MetaBrain AFR subset (n=311). We used the expression matrix as described for the gene set enrichment analysis and prepared the gene expression for AFR in an identical manner. We did not run PICALO on the AFR samples, but rather we first calculated the Pearson correlation between PIC values and gene expression levels in the EUR subset. In the AFR subset, we then used a dot product to associate the AFR gene expression levels with PIC-gene correlations observed in the EUR subset. This provides an estimate of the EUR PIC values in the AFR subset. We then performed ieQTL analysis in the EUR and AFR datasets as outlined in ‘Detection of eQTL interacting with PICs / expression PCs’ but in this case we did not perform conditional analysis. Per PIC we calculated a BH-FDR over the p-values of the ieQTLs that were significant in EUR.

Furthermore, we calculated three different measurements of agreement (allelic concordance, π ^29^ and R ^30^) using the ieQTLs that were significant in EUR. To calculate the π_1_ we took the AFR ieQTL p-38 values and calculated the proportion of true null p-values (π_0_) with the pi0est function from qvalue and subsequently calculated π_1_ as 1 – π_0_ . Similarly, we calculated the R_b_ by comparing the interaction beta and standard errors between EUR and AFR.

## Data availability and accession codes

The raw RNA-seq data of BIOS can be obtained from the European Genome-phenome Archive (EGA; accession EGAS00001001077). Genotype data are available from the respective biobanks: LLS (http://www.leidenlangleven.nl/en/home), LifeLines (https://lifelines.nl/lifelines-research/access-to-lifelines), CODAM and RS (http://www.erasmusmc.nl/epi/research/The-Rotterdam-Study/?lang=en). The MetaBrain dataset is comprised of previously published human brain eQTL datasets. The majority of the used datasets in this paper are available upon request, or through online repositories after signing data access agreements. We have listed the mode of access for each of the included datasets below.

TargetALS^39^ TargetALS data was pushed directly from the NY Genome center to our sftp server. CMC^40^ CMC data was downloaded from https://www.synapse.org/ using synapse client (https://python-docs.synapse.org/build/html/index.html). Accession code: syn2759792.

GTEx^41^ GTEx was downloaded from SRA using fastq-dump of the SRA toolkit (http://www.ncbi.nlm.nih.gov/Traces/sra/sra.cgi?cmd=show&f=software&m=software&s=software). Access has been requested and granted through dbGaP. Accession code: phs000424.v7.p2 AMP-AD^42^ AMP-AD data has been downloaded from synapse. Accession code: syn2580853. snRNA- seq was collected using Synapse accession code: syn18485175.

ENA^43^ ENA data has been downloaded from the European Nucleotide Archive. The identifiers of the 76 included studies and 2021 brain samples are listed in de Klein et al. 2021.

CMC_HBCC^40^ CMC_HBCC data was downloaded from https://www.synapse.org/ using synapse client (https://python-docs.synapse.org/build/html/index.html). Accession code: syn10623034

BrainSeq^44^ BrainSeq data was downloaded from https://www.synapse.org/ using synapse client (https://python-docs.synapse.org/build/html/index.html). Accession code: syn12299750 UCLA_ASD^45^ UCLA_ASD data was downloaded from https://www.synapse.org/using synapse client (https://python-docs.synapse.org/build/html/index.html). Accession code: syn4587609 BrainGVEx^45^ BrainGVEx data was downloaded from https://www.synapse.org/ using synapse client (https://python-docs.synapse.org/build/html/index.html). Accession code: syn4590909

BipSeq^45^ BipSeq data was downloaded from https://www.synapse.org/ using synapse client (https://python-docs.synapse.org/build/html/index.html). Accession code: syn5844980 NABEC^46^ NABEC data was downloaded from dbgap. Accession code: phs001301.v1.p1 BLUEPRINT^36^ expression data was downloaded from http://dcc.blueprint-epigenome.eu

## Ethical compliance

All cohorts included in this study enrolled participants with informed consent, and collected and analyzed data in accordance with ethical and institutional regulations. The information about individual institutional review board approvals is available in the original publications for each cohort. Where applicable, data access agreements were signed by the investigators prior to acquisition of the data, either to the UMCG (AMP-AD, CMC, GTEx, CMC_HBCC, BrainSeq, UCLA_ASD, BrainGVEX, BipSeq, NABEC), or Biogen (TargetALS) which state the data usage terms. To protect the privacy of the participants, data access was restricted to the investigators of this study, as defined in those data access agreements. Per data use agreements, only summary level data is made publicly available and strictly mentioned in the disclaimer that they cannot be used to re-identify study participants.

## Code availability

The PICALO software, as well as custom scripts used in this manuscript, are available at https://github.com/molgenis/PICALO.

## Acknowledgments

We thank the donors of the brain tissues underlying the RNA-seq data used for this study and their families for their willingness to donate samples for research. This study makes use of data generated by the Biobank-based Integrative Omics Study. A full list of the investigators is available from http://www.bbmri.nl/en-gb/activities/rainbow-projects/bios. Funding for the project was provided by the Netherlands Organization for Scientific Research under award number 184021007, dated July 9, 2009 and made available as a Rainbow Project of the Biobanking and Biomolecular Research Infrastructure Netherlands (BBMRI-NL). Samples were contributed by the UMCG ‘LifeLines’ collection^2^ (http://lifelines.nl), the LUMC Longevity Study^47^ (http://www.lumc.nl/con/2095/83047/86636/86648/), the Rotterdam Study^48^ (http://www.epib.nl/research/ergo.html), the VU Netherlands Twin Register^49^ (www.tweelingenregister.org), Cohort study on Diabetes and Artherosclerosis (CODAM) ^50^, and Prospective ALS Study Netherlands (PAN)^51^. A small subset of samples was also full genome sequenced within the GoNL project^52^. We acknowledge the contributions of the investigators to this study (Supplementary Note). We would like to thank the Center for Information Technology of the University of Groningen for their support and for providing access to the Peregrine high-performance computing cluster, as well as the UMCG Genomics Coordination center, the UG Center for Information Technology and their sponsors BBMRI-NL and TarGet for storage and compute infrastructure. We thank Urmo Võsa for the script he provided to calculate R_b_ statistics. Finally, we thank Carlos Urzúa-Traslaviña and Kate McIntyre for the editorial assistance.

## ROSMAP

The results published here are in whole or in part based on data obtained from the AMP-AD Knowledge Portal (<u>doi:10.7303/syn2580853</u>) Study data were provided by the Rush Alzheimer’s Disease Center, Rush University Medical Center, Chicago. Data collection was supported through funding by NIA grants P30AG10161, R01AG15819, R01AG17917, R01AG30146, R01AG36836, U01AG32984, U01AG46152, the Illinois Department of Public Health, and the Translational Genomics Research Institute. Genotype data: doi:10.1038/mp.2017.20. RNAseq: <u>doi:10.1038/s41593-018-0154-9</u>. snRNA-seq: <u>doi:10.7303/syn18485175</u>

## Mayo

The results published here are in whole or in part based on data obtained from the AMP-AD Knowledge Portal (<u>doi:10.7303/syn2580853</u>). Study data were provided by the following sources: The Mayo Clinic Alzheimer’s Disease Genetic Studies, led by Dr. Nilufer Taner and Dr. Steven G. Younkin, Mayo Clinic, Jacksonville, FL using samples from the Mayo Clinic Study of Aging, the Mayo Clinic Alzheimer’s Disease Research Center, and the Mayo Clinic Brain Bank. Data collection was supported through funding by NIA grants P50 AG016574, R01 AG032990, U01 AG046139, R01 AG018023, U01 AG006576, U01 AG006786, R01 AG025711, R01 AG017216, R01 AG003949, NINDS grant R01 NS080820, CurePSP Foundation, and support from Mayo Foundation. Study data includes samples collected through the Sun Health Research Institute Brain and Body Donation Program of Sun City, Arizona. The Brain and Body Donation Program is supported by the National Institute of Neurological Disorders and Stroke (U24 NS072026 National Brain and Tissue Resource for Parkinsons Disease and Related Disorders), the National Institute on Aging (P30 AG19610 Arizona Alzheimer’s Disease Core Center), the Arizona Department of Health Services (contract 211002, Arizona Alzheimer’s Research Center), the Arizona Biomedical Research Commission (contracts 4001, 0011, 05-901 and 1001 to the Arizona Parkinson’s Disease Consortium) and the Michael J. Fox Foundation for Parkinson’s Research. <u>doi:10.1038/sdata.2016.89</u>

## MSBB

The results published here are in whole or in part based on data obtained from the AMP-AD Knowledge Portal (<u>doi:10.7303/syn2580853</u>). These data were generated from postmortem brain tissue collected through the Mount Sinai VA Medical Center Brain Bank and were provided by Dr. Eric Schadt from Mount Sinai School of Medicine.

## CMC

Data were generated as part of the CommonMind Consortium supported by funding from Takeda Pharmaceuticals Company Limited, F. Hoffman-La Roche Ltd and NIH grants R01MH085542, R01MH093725, P50MH066392, P50MH080405, R01MH097276, RO1-MH-075916, P50M096891, P50MH084053S1, R37MH057881, AG02219, AG05138, MH06692, R01MH110921, R01MH109677, R01MH109897, U01MH103392, and contract HHSN271201300031C through IRP NIMH. Brain tissue for the study was obtained from the following brain bank collections: The Mount Sinai NIH Brain and Tissue Repository, the University of Pennsylvania Alzheimer’s Disease Core Center, the University of Pittsburgh NeuroBioBank and Brain and Tissue Repositories, and the NIMH Human Brain Collection Core. CMC Leadership: Panos Roussos, Joseph Buxbaum, Andrew Chess, Schahram Akbarian, Vahram Haroutunian (Icahn School of Medicine at Mount Sinai), Bernie Devlin, David Lewis (University of Pittsburgh), Raquel Gur, Chang-Gyu Hahn (University of Pennsylvania), Enrico Domenici (University of Trento), Mette A. Peters, Solveig Sieberts (Sage Bionetworks), Thomas Lehner, Stefano Marenco, Barbara K. Lipska (NIMH).

## GTEx

The Genotype-Tissue Expression (GTEx) Project was supported by the Common Fund of the Office of the Director of the National Institutes of Health (commonfund.nih.gov/GTEx). Additional funds were provided by the NCI, NHGRI, NHLBI, NIDA, NIMH, and NINDS. Donors were enrolled at Biospecimen Source Sites funded by NCI\Leidos Biomedical Research, Inc. subcontracts to the National Disease Research Interchange (10XS170), Roswell Park Cancer Institute (10XS171), and Science Care, Inc. (X10S172). The Laboratory, Data Analysis, and Coordinating Center (LDACC) was funded through a contract (HHSN268201000029C) to the The Broad Institute, Inc. Biorepository operations were funded through a Leidos Biomedical Research, Inc. subcontract to Van Andel Research Institute (10ST1035). Additional data repository and project management were provided by Leidos Biomedical Research, Inc.(HHSN261200800001E). The Brain Bank was supported supplements to University of Miami grant DA006227. Statistical Methods development grants were made to the University of Geneva (MH090941 & MH101814), the University of Chicago (MH090951, MH090937, MH101825, & MH101820), the University of North Carolina - Chapel Hill (MH090936), North Carolina State University (MH101819), Harvard University (MH090948), Stanford University (MH101782), Washington University (MH101810), and to the University of Pennsylvania (MH101822). The datasets used for the analyses described in this manuscript were obtained from dbGaP at http://www.ncbi.nlm.nih.gov/gap through dbGaP accession number phs000424.v7.p2 on 02/27/2020.

## NABEC

Data was collected from dbGAP accession phs001301.v1.p1, which was generated by J. R. Gibbs, M. van der Brug, D. Hernandez, B. Traynor, M. Nalls, S-L. Lai, S. Arepalli, A. Dillman, I. Rafferty, J. Troncoso, R. Johnson, H. R. Zielke, L. Ferrucci, D. Longo, M.R. Cookson, and A.B. Singleton. The NABEC dataset was generated at National Institute on Aging, Bethesda, MD, USA, Institute of Neurology, University College London, London, UK, The Scripps Research Institute, Jupiter, FL, USA, Johns Hopkins University, Baltimore, MD, USA, and the University of Maryland Medical School, Baltimore, MD, USA. NABEC was funded by Z01 AG000949-02. National Institutes of Health, Bethesda, MD, USA and Z01 AG000015-49. National Institutes of Health, Bethesda, MD, USA.

## TargetALS

This data set was generated and supported by the following: Target ALS Human Postmortem Tissue Core, New York Genome Center for Genomics of Neurodegenerative Disease, Amyotrophic Lateral Sclerosis Association and TOW Foundation.

## European Nucleotide Archive

We would like to thank all donors and their families, principal investigators and their funding bodies for each of the projects included from the European Nucleotide Archive.

## UCLA ASD, Bipseq, BrainGVEx and LIBD

Data were generated as part of the PsychENCODE Consortium supported by: U01MH103339, U01MH103365, U01MH103392, U01MH103340, U01MH103346, R01MH105472, R01MH094714, R01MH105898, R21MH102791, R21MH105881, R21MH103877, and P50MH106934 awarded to: Schahram Akbarian (Icahn School of Medicine at Mount Sinai), Gregory Crawford (Duke), Stella Dracheva (Icahn School of Medicine at Mount Sinai), Peggy Farnham (USC), Mark Gerstein (Yale), Daniel Geschwind (UCLA), Thomas M. Hyde (LIBD), Andrew Jaffe (LIBD), James A. Knowles (USC), Chunyu Liu (UIC), Dalila Pinto (Icahn School of Medicine at Mount Sinai), Nenad Sestan (Yale), Pamela Sklar (Icahn School of Medicine at Mount Sinai), Matthew State (UCSF), Patrick Sullivan (UNC), Flora Vaccarino (Yale), Sherman Weissman (Yale), Kevin White (UChicago) and Peter Zandi (JHU).

## Lifelines

The LifeLines Cohort Study (http://www.lifelines.nl), and generation and management of GWAS genotype data for it, is supported by the Netherlands Organization of Scientific Research (NWO, grant 175.010.2007.006), the Dutch government’s Economic Structure Enhancing Fund (FES), the Ministry of Economic Affairs, the Ministry of Education, Culture and Science, the Ministry for Health, Welfare and Sports, the Northern Netherlands Collaboration of Provinces (SNN), the Province of Groningen, the University Medical Center Groningen, the University of Groningen, the Dutch Kidney Foundation and Dutch Diabetes Research Foundation.

## Rotterdam Study

The Rotterdam Study is funded by Erasmus Medical Center and Erasmus University, Rotterdam, Netherlands Organization for the Health Research and Development (ZonMw), the Research Institute for Diseases in the Elderly (RIDE), the Ministry of Education, Culture and Science, the Ministry for Health, Welfare and Sports, the European Commission (DG XII), and the Municipality of Rotterdam. The authors are grateful to the study participants, the staff from the Rotterdam Study and the participating general practitioners and pharmacists. The generation and management of GWAS genotype data for the Rotterdam Study is supported by the Netherlands Organisation of Scientific Research NWO Investments (nr. 175.010.2005.011, 911-03-012). This study is funded by the Netherlands Genomics Initiative (NGI)/Netherlands Organisation for Scientific Research (NWO) project nr. 050-060-810.

## LUMC Longevity Study

The LUMC Longevity Study was supported by a grant from the Innovation-Oriented Research Program on Genomics (SenterNovem IGE01014 and IGE05007), the Centre for Medical Systems Biology and the National Institute for Healthy Ageing (Grant 05040202 and 05060810), all in the framework of the Netherlands Genomics Initiative/Netherlands Organization for Scientific Research.;

## VU Netherlands Twin Register

For sponsorship of the VU Netherlands Twin Register please refer to www.tweelingenregister.org. Cohort study on Diabetes and Atherosclerosis Maastricht (CODAM)

The initiation of the Cohort on Diabetes and Atherosclerosis Maastricht (CODAM) was supported by grants from the Netherlands Organization for Scientific Research (NWO grant 940–35–034), the Dutch Diabetes Research Foundation (DFN 98.901). Generation of GWAS genotype data for CODAM was supported by Netherlands Organization for Scientific Research (NWO 184.021.007) and made available as a Complementation Project of the Biobanking and Biomolecular Research Infrastructure Netherlands (BBMRI-NL).

## Prospective ALS Study Netherlands (PAN)

The PAN study (Prospective Study of ALS in the Netherlands) is funded by the Stichting ALS (www.als-stichting.nl)

## BLUEPRINT

This study makes use of data generated by the BLUEPRINT Consortium (release 8). A full list of the investigators who contributed to the generation of the data is available from www.blueprint-epigenome.eu. Funding for the project was provided by the European Union’s Seventh Framework Programme (FP7/2007-2013) under grant agreement no 282510 – BLUEPRINT.

## Author contributions

- Conceptualization: L.F.
- Methodology: M.V. and L.F.
- Software: M.V., P.D., and B.V.
- Validation: M.V.
- Formal Analysis: M.V., P.D., and B.V.
- Investigation: M.V., P.D., B.V., S.A.S., J.F., and A.Z.
- Resources: E.A.T. and H.R.
- Data curation: M.V.
- Writing – original draft: M.V. and P.D.
- Writing – review & editing: P.D., H.W., and L.F.
- Visualization: M.V., P.D., and H.W.
- Supervision: P.D., H.W., and L.F.
- Funding acquisition: P.D., E.A.T., H.R., L.F.

Roles as defined by: CRediT (Contributor Roles Taxonomy)

## Supplementary data

**Fig. S1.**
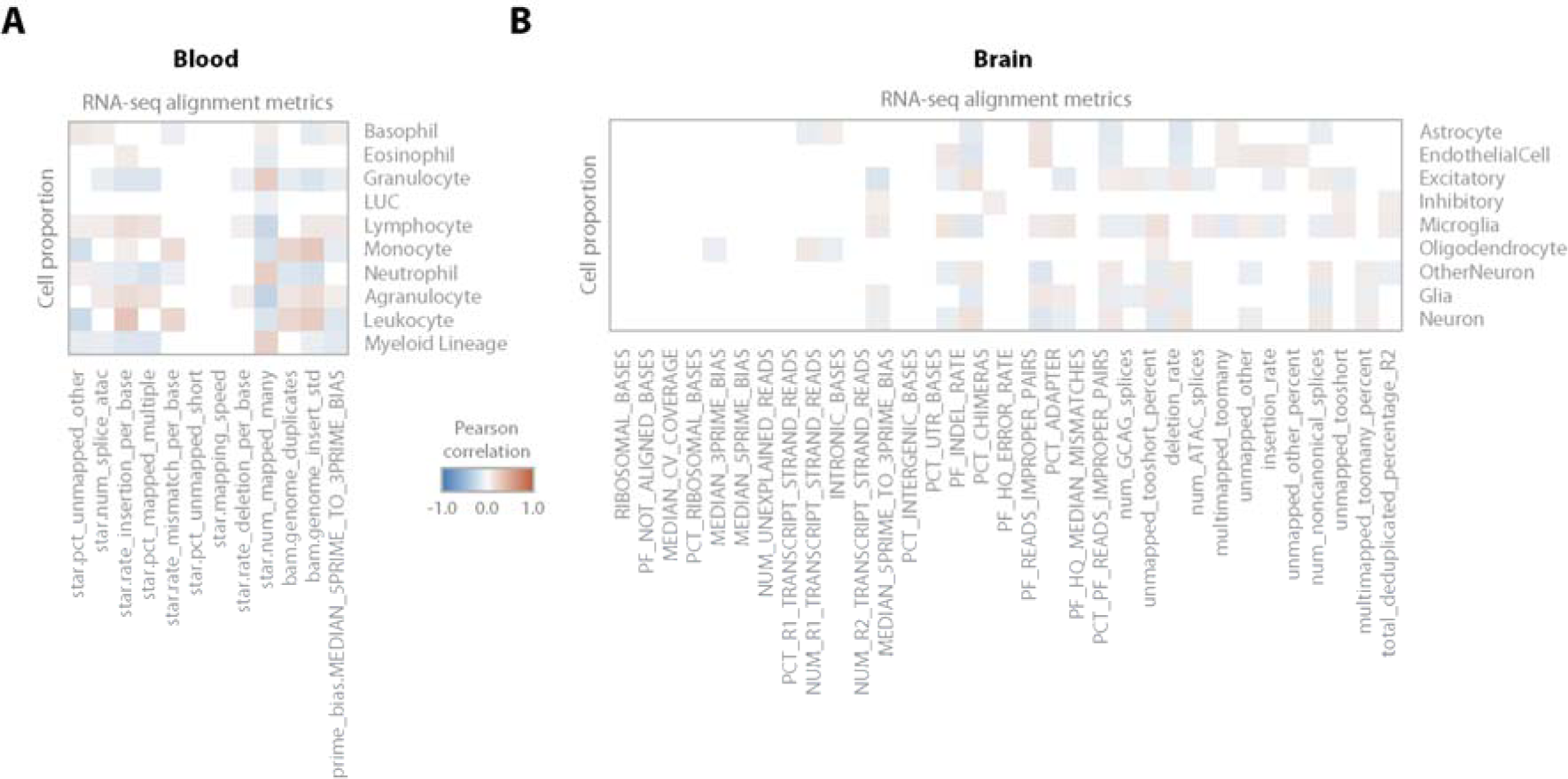
Correlation between RNA-seq alignment metrics and cell type proportions. *Pearson correlations between RNA-seq alignment metrics and measured cell type proportions in blood (A), and predicted cell type proportions in brain (B). The correlations in brain are less evident, most likely because RNA- seq alignment metrics were explicitly corrected for prior to predicting cell proportions in this dataset*.

**Fig. S2.**
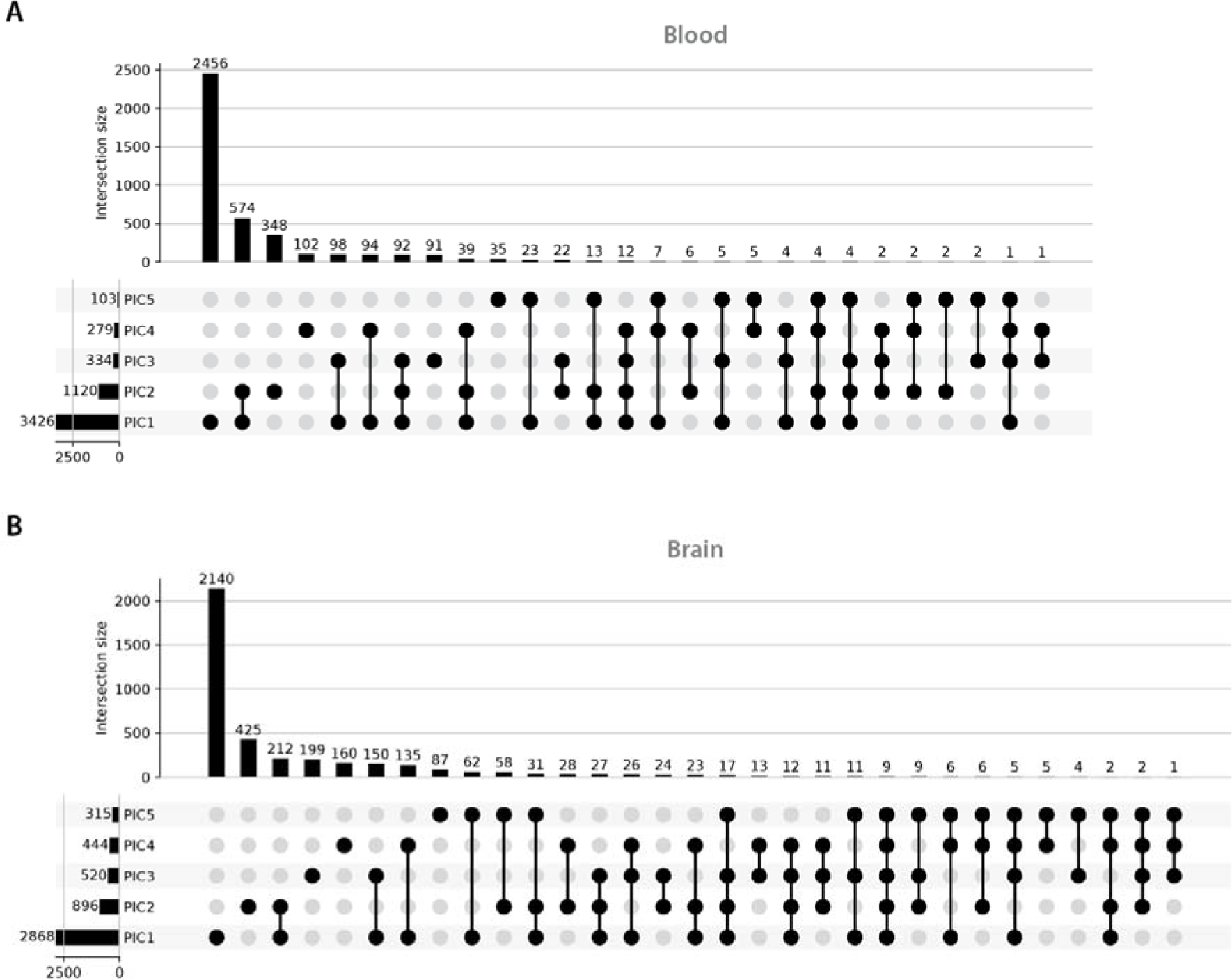
Unique and overlapping PIC interaction eQTLs. *Unique and overlapping PIC ieQTLs for the first five PICs in blood (A) and brain (B). The majority of eQTL interacted with a single PIC*.

**Fig. S3.**
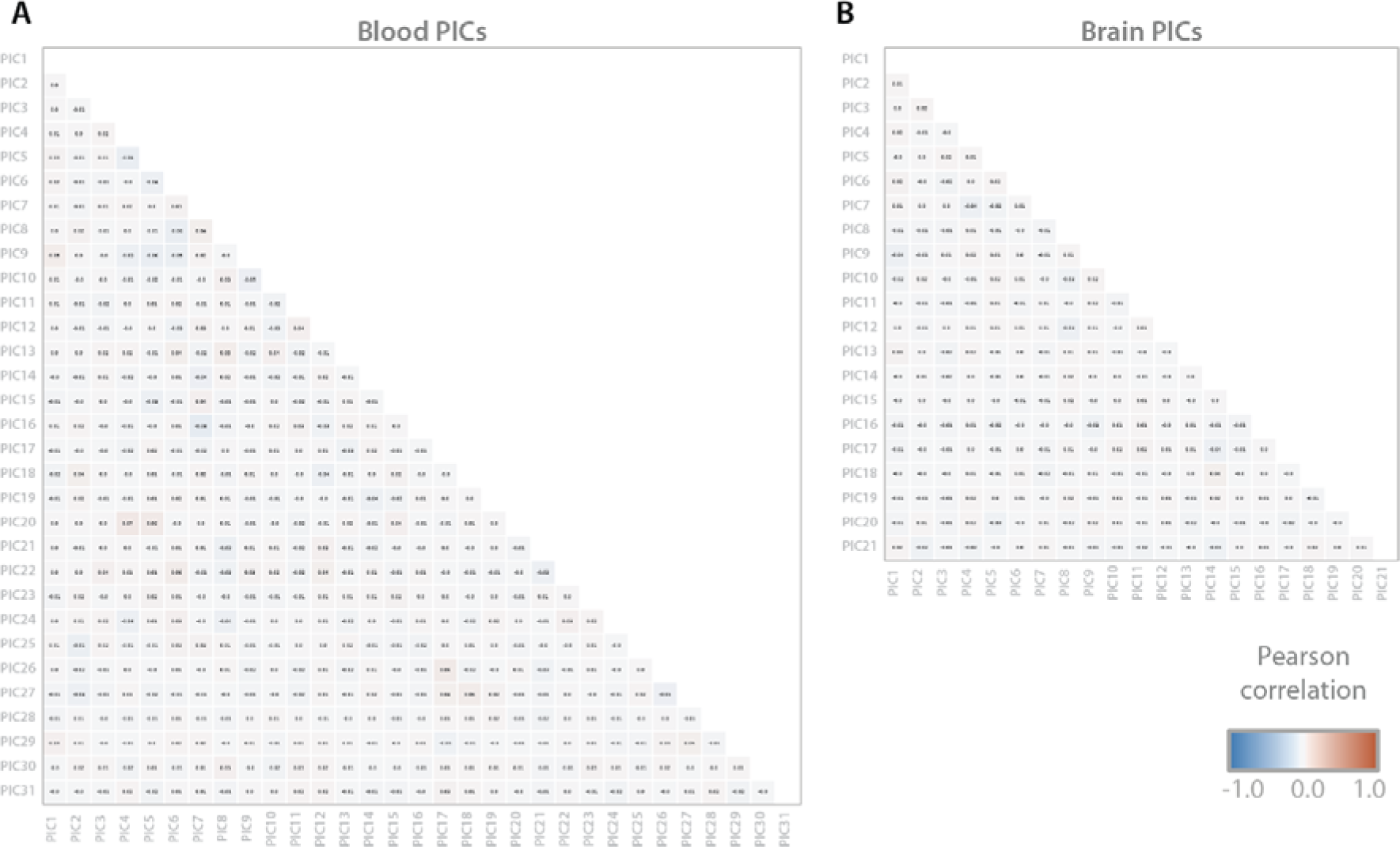
Correlation between PICs. Pearson correlation between identified PICs in blood (A) and brain (B)*. No correlation (max Pearson r*≤*0.07) between PICs is found in either dataset*.

**Fig. S4.**
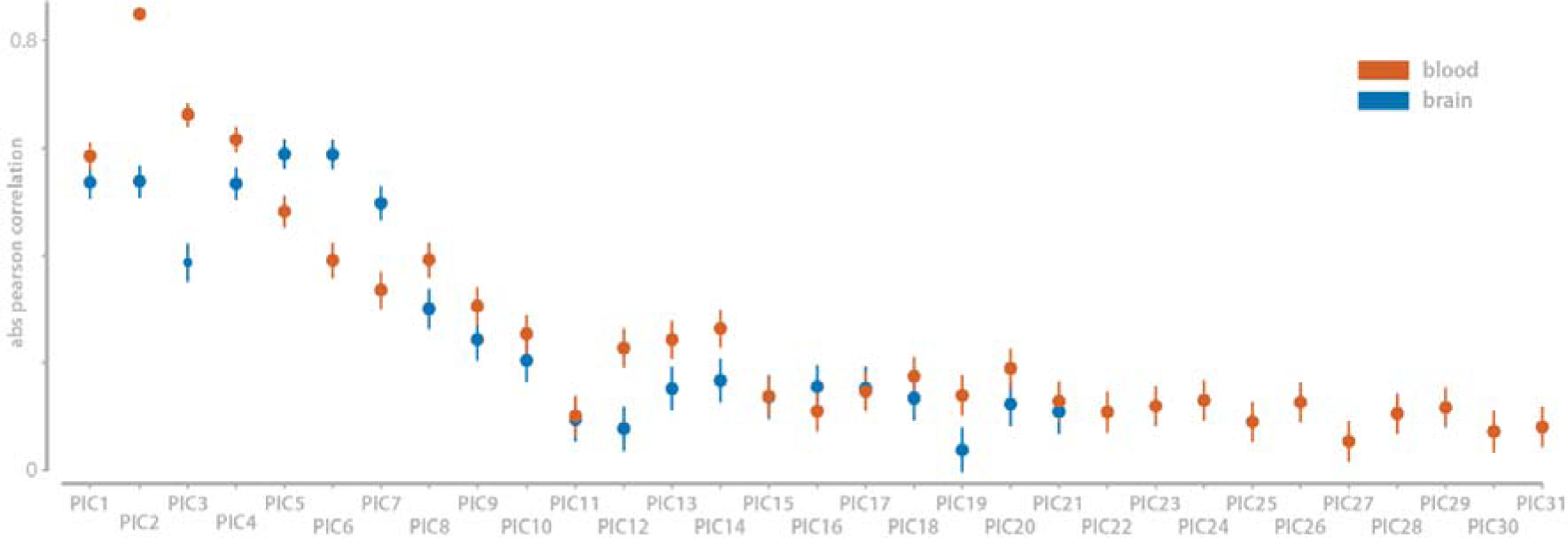
Correlation between PIC and initial guess. The Pearson correlation between the initial guess (expression PC; before optimization) and the resulting PIC (after optimization) for blood and brain. The error bars indicate the 95% confidence interval. The correlation before and after optimization decreases as the PICs capture less interaction variance.

**Fig. S5.**
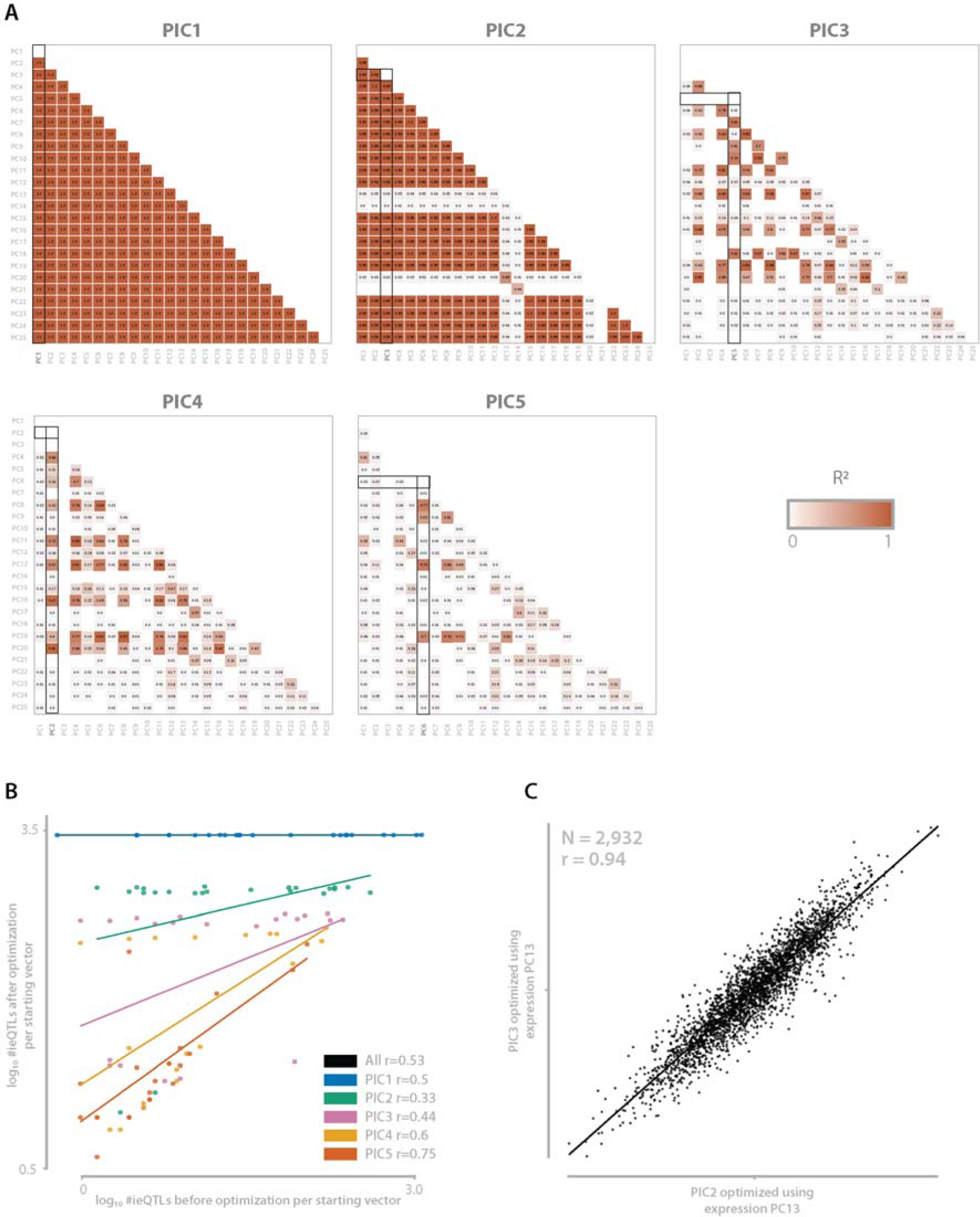
Correlation between identified PICs using different initial guesses. *(A) Comparison of PICs using different expression PCs as initial guess. The first five PICs are tested, for each PIC the initial guess (i.e. expression PC) with the highest number of ieQTLs before optimization is used as PIC and is corrected for before analyzing PIC2 (initial guesses used PIC1=PC1, PIC2=PC3, PIC3=PC5, PIC4=PC2, PIC5=PC6). For each of initial guesses the resulting outcome is compared to all the other outcomes when using different starting vectors. Each cell contains the R^2^ of these comparisons. Only correlations with a BH-FDR≤0.05 are shown. PIC1 for example, shows that regardless of which initial guess used for optimization, the resulting outcome is the same. (B) Visualization of the relationship between the number of identified ieQTLS before (x- axis) and after (y-axis) optimization. For the top PICs there is no relation between the number of ieQTLs before and after optimization. However, this dependence increases as the proportion of interaction variance the PICs capture decreases. (C) Comparison of PIC2 optimized using expression PC13 as initial guess versus PIC3 optimized using expression PC13. Note that the PICALO outcome for PIC2 using expression PC13 was not removed when accessing PIC3 since a different PC had more ieQTLs prior to optimization. The PIC2-PC13 outcome, which is likely a local minimum, can therefore be reidentified when optimizing expression PC13 for PIC3, highlighting the robustness of the method*.

**Fig. S6.**
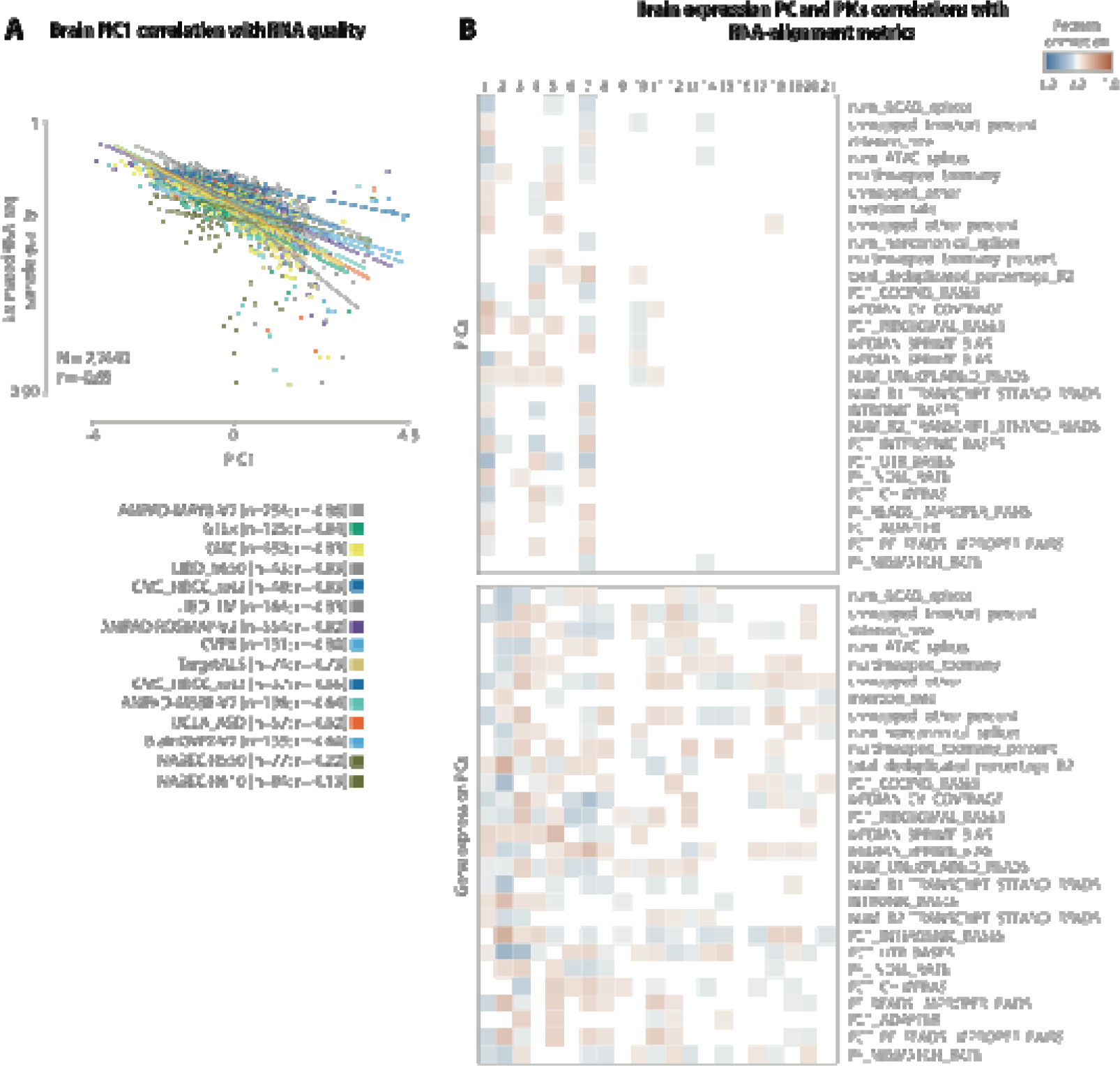
Brain PIC and expression PC correlations with estimated RNA-seq sample quality and RNA-seq alignment metrics. *(A) Regression plot showing the correlation between PIC1 and estimated RNA-seq sample quality calculated as the per sample expression correlation with the overall average expression in brain. (B) Pearson correlation heatmaps correlating PICs (top) as well as expression PC (bottom) to RNA-seq alignment metrics in brain. The correlation p-values are corrected for multiple testing with Benjamini-Hochberg and only correlations with an FDR≤0.05 are shown. Note that many of the expression PCs correlate significantly with RNA-seq alignment metrics while only a limited number of PICs show significant correlations*.

**Fig. S7.**
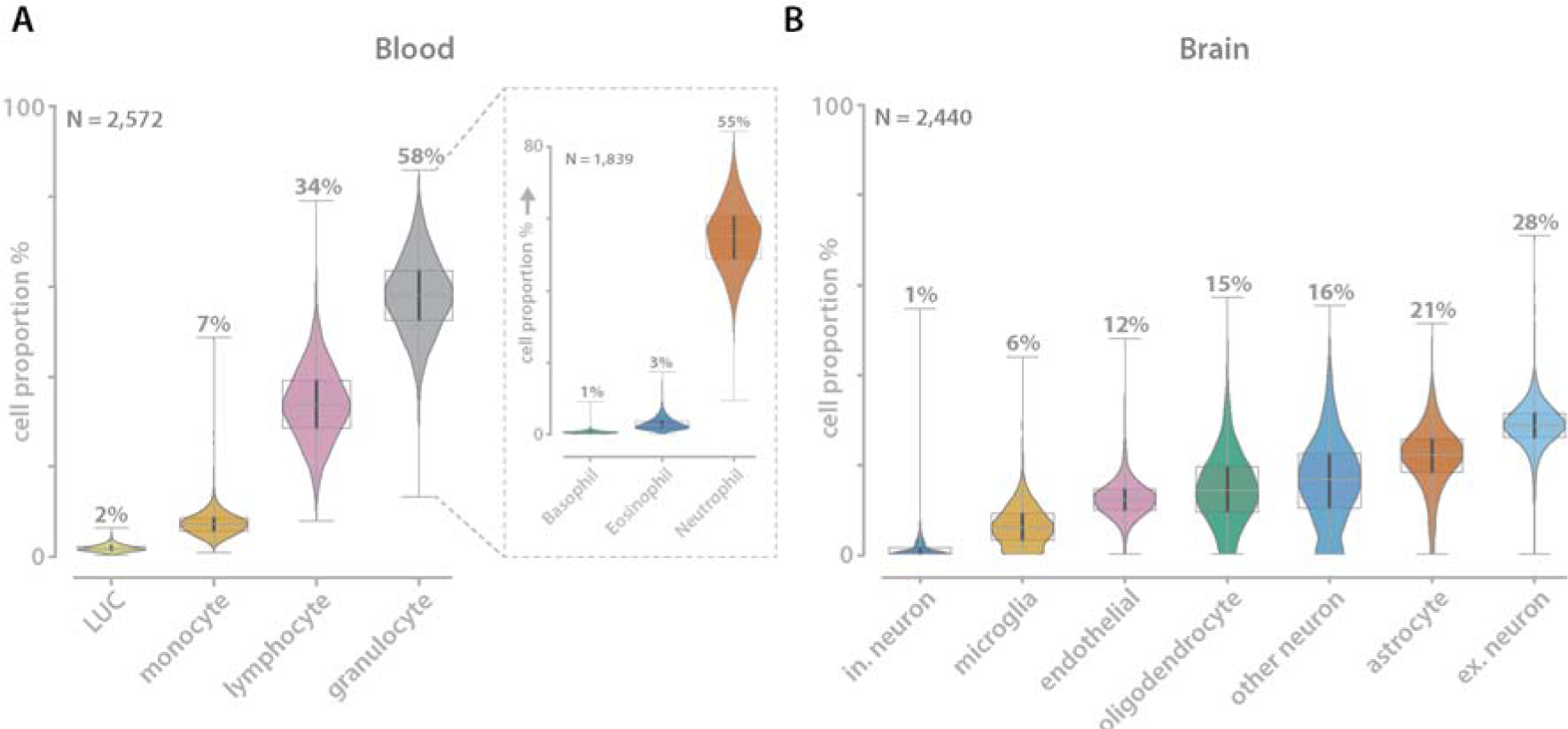
Cell type proportion in blood (measured) and brain (predicted) *(A) Measured cell type proportions for the blood samples. For a subset of samples (n=1,839) the granulocytes are further distinguished into sub cell types. (B) Predicted cell type proportions for the brain samples as described by de Klein et al. 2021*.

**Fig. S8.**
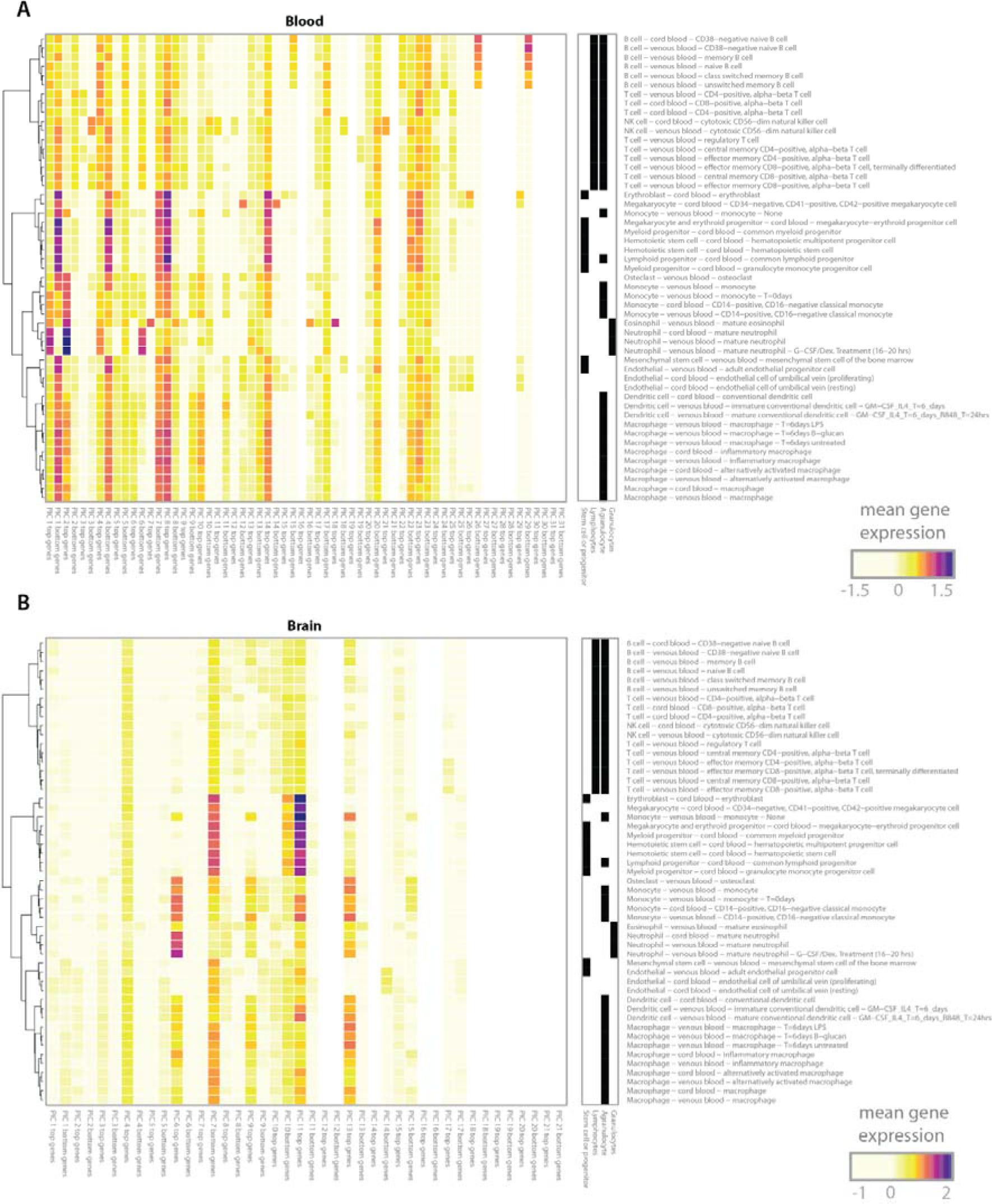
Gene set enrichment in BLUEPRINT data. *Mean gene expression of purified BLUEPRINT RNA-seq expression for the top 200 positively and top 200 negatively correlated genes in blood (A) and brain (B). The FPKM+1 BLUEPRINT expression data is log2 transformed and center and scaled per sample*.

**Fig. S9.**
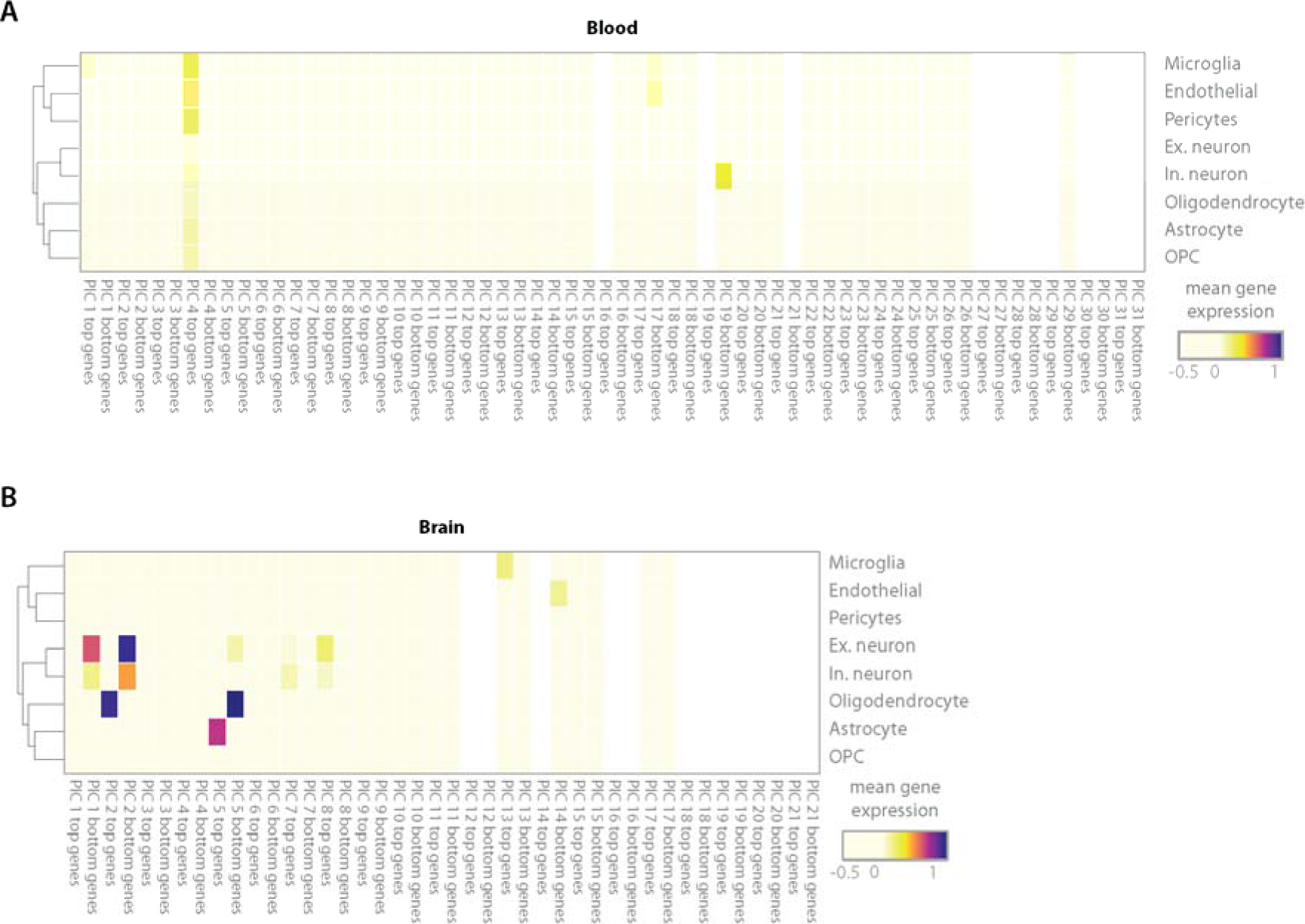
Brain gene set enrichment in ROSMAP single-nucleus data. Mean gene expression of the ROSMAP single-nucleus RNA-seq expression for top 200 positively and top 200 negatively correlated genes in blood (A) and brain (B). The ROSMAP expression data is log_2_ transformed and center and scaled per sample.

**Fig. S10.**
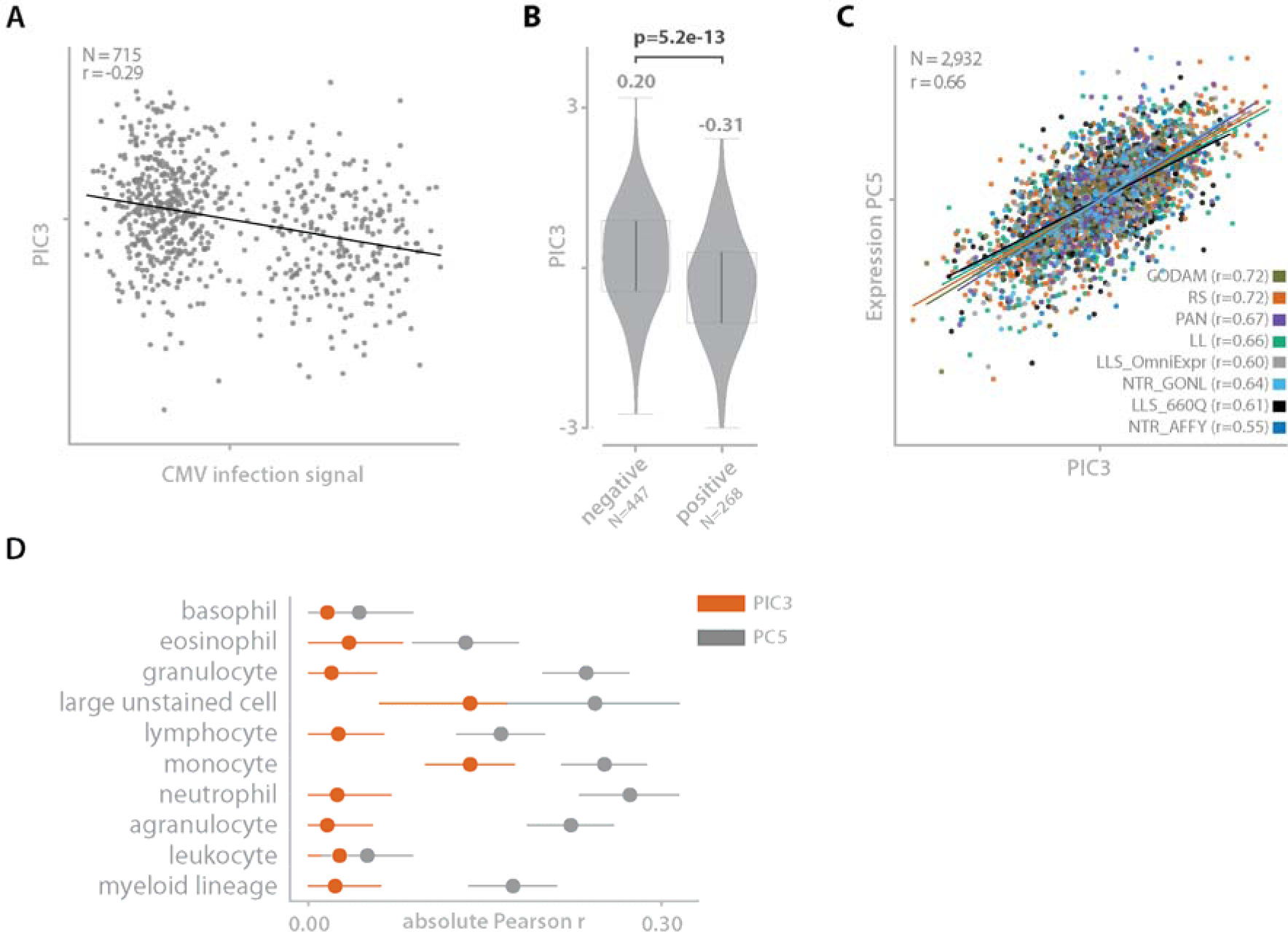
PIC3 correlated with antibody PC describing prior CMV infection. (A) Regression plot comparing the PIC3 to CMV infection signal (aggregate of multiple lgG antibody profiles) for the LifeLines (LL) cohort. (B) Violin plot comparing the PIC3 scores for samples with a negative or a positive CMV infection signal. The significance between the two groups is calculated using a two-sided Mann–Whitney U test. (C) Comparison of PIC3 and expression PC5, both correlating with CMV infection signal. (D) Forest plot showing the absolute Pearson correlations and 95% confidence intervals for PIC3 and expression PC5 compared to cell type proportions. PIC3 shows very low correlations while expression PC5 shows higher correlation, suggesting that PIC3 interactions are less biased by cell type confounding.

**Fig. S11.**
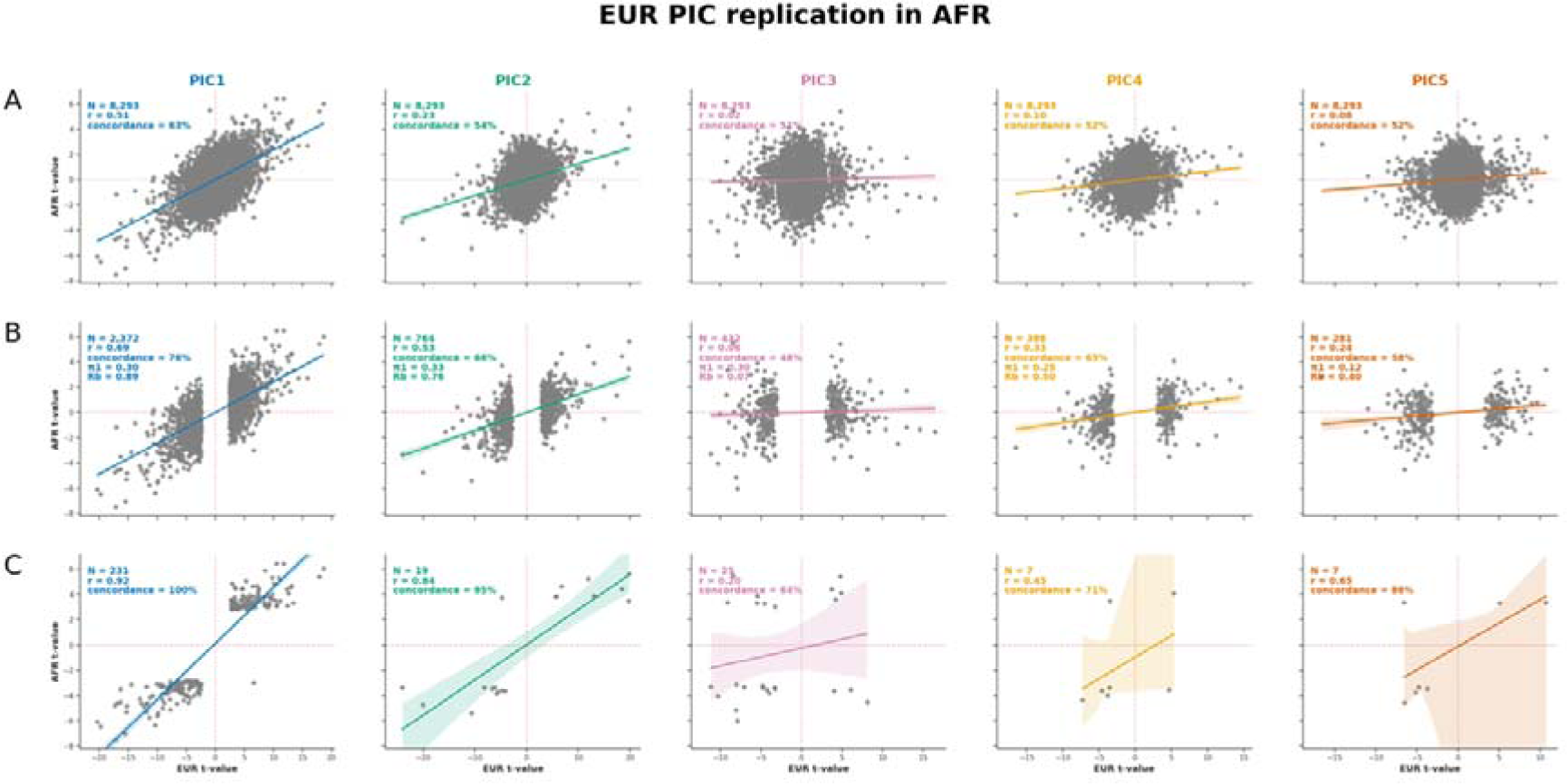
Brain PICs replication in independent dataset. *Replication of MetaBrain EUR interaction eQTLs identified in MetaBrain AFR independent dataset. Each figure in this plot represents a comparison between MetaBrain EUR (x-axis) and MetaBrain AFR (y-axis). Each dot represents one ieQTL, and the legend shows the sample size, Pearson correlation coefficient, the allelic concordance, and, if applicable, the R_b_ and* π*1 statistics. Each column is a comparison between equivalent cell types in both datasets. Each row illustrates a different filtering on which eQTLs are shown. The x-axis always denotes the interaction t-value in EUR, the y-axis always denotes the interaction t-value in AFR. (A) All overlapping ieQTLs (B) ieQTLs filtered on being significant in EUR (C) ieQTLs filtered on being significant in both datasets*.

**Fig. S12.**
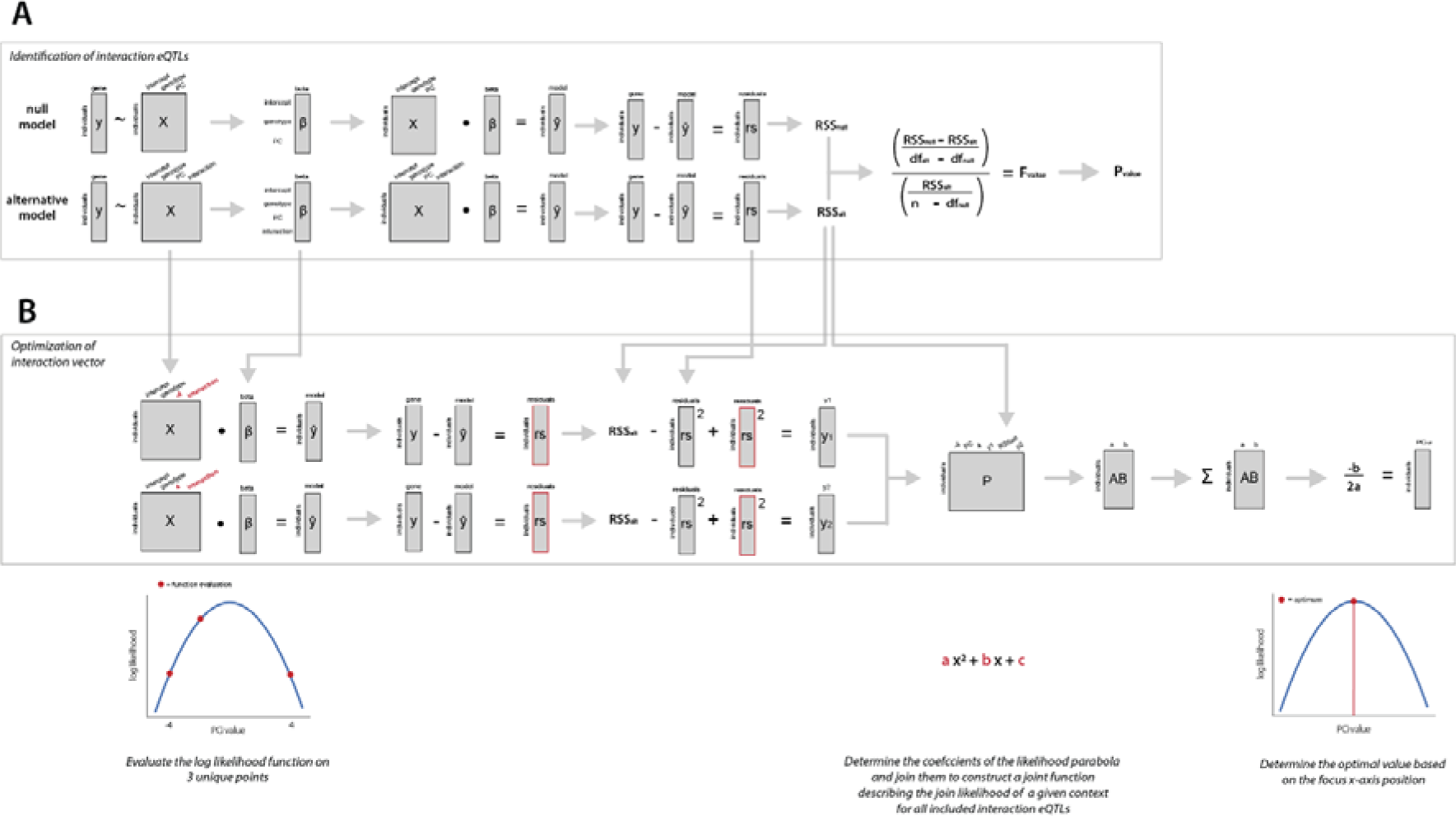
PICALO interaction eQTL identification and optimization. *Interaction eQTL identification and optimization steps as implemented in PICALO. (A) the gene expression is modelled using two models; one without an interaction term (null model) and one with an interaction term (alternative model). By comparing the residual sum of squares of both models the significance of the interaction term can be determined using a F-test. (B) For the eQTLs that have a significant interaction, the optimal context value is determined per sample. For this the residuals are minimized over all included ieQTL per sample. Per sample and per ieQTL the alternative model is re-evaluated using -4 and +4 as context value. This, together with the initial guess, gives three coordinates used to determine the unique second-degree polynomial that intersects these coordinates. The coefficients of the parabola are then summed together per sample over all ieQTLs to construct a combined function describing the joint log-likelihood. The optimal context value is then determined by calculating the focus of the combined function*.

**Fig. S13.**
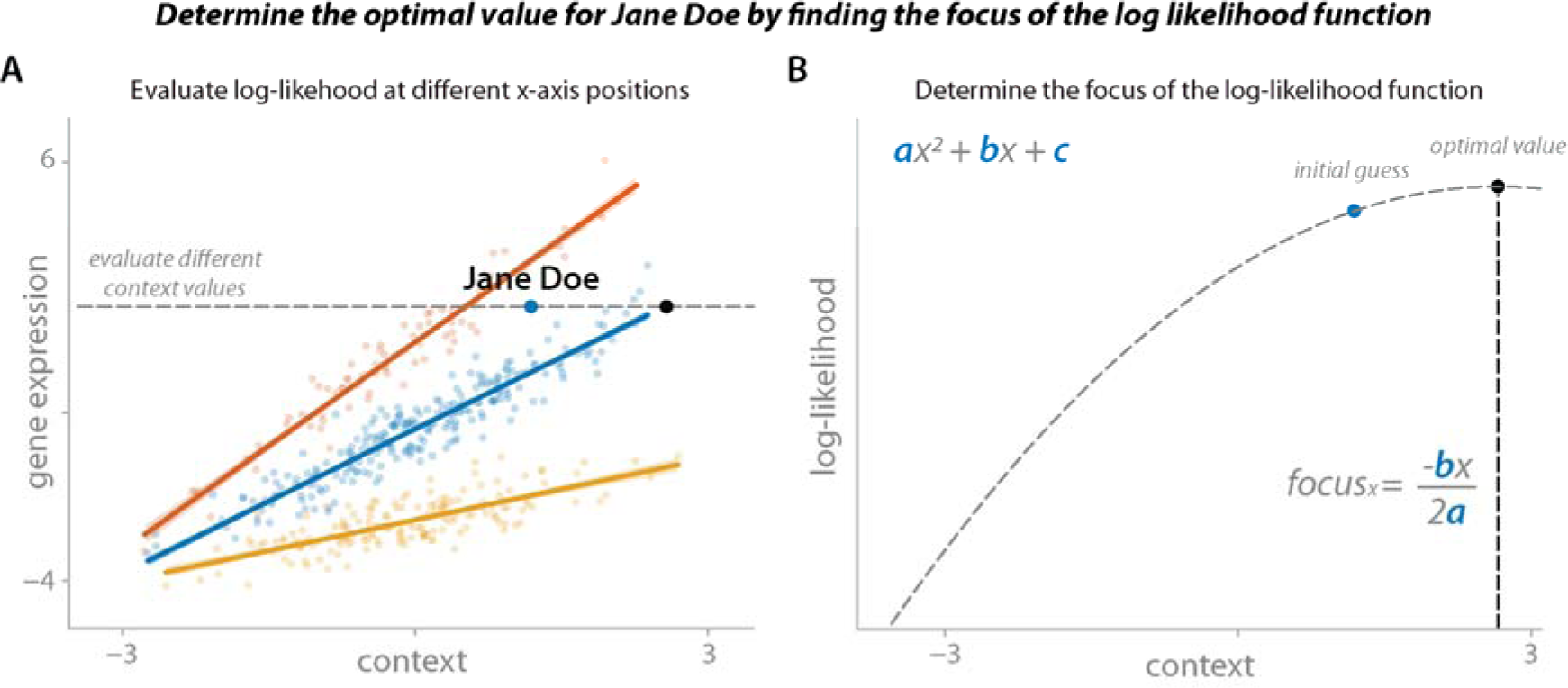
Single interaction eQTL optimization example. *Cartoon example showing how the optimal context value for a single sample (Jane Doe) is determined when applying PICALO to a single ieQTL. (A) The ieQTL model is fitted over all samples. Per sample, the context value is adjusted (moved along the x-axis) and the change in log-likelihood is evaluated. Note that all other parameters, including the interaction beta, are not updated during this process. (B). The change in log-likelihood follows a second-degree polynomial function, the focus of which gives the context value with the maximum log-likelihood. To illustrate: in the case of a single ieQTL, this translates to sliding each sample in turn along the x-axis until it intersects with the regression line of its genotype group. In the case of multiple ieQTLs being optimized simultaneously (as is the case in PICALO), the log-likelihood functions are summed together and the optimal context value is determined over the joint log-likelihood*.

**Fig. S14.**
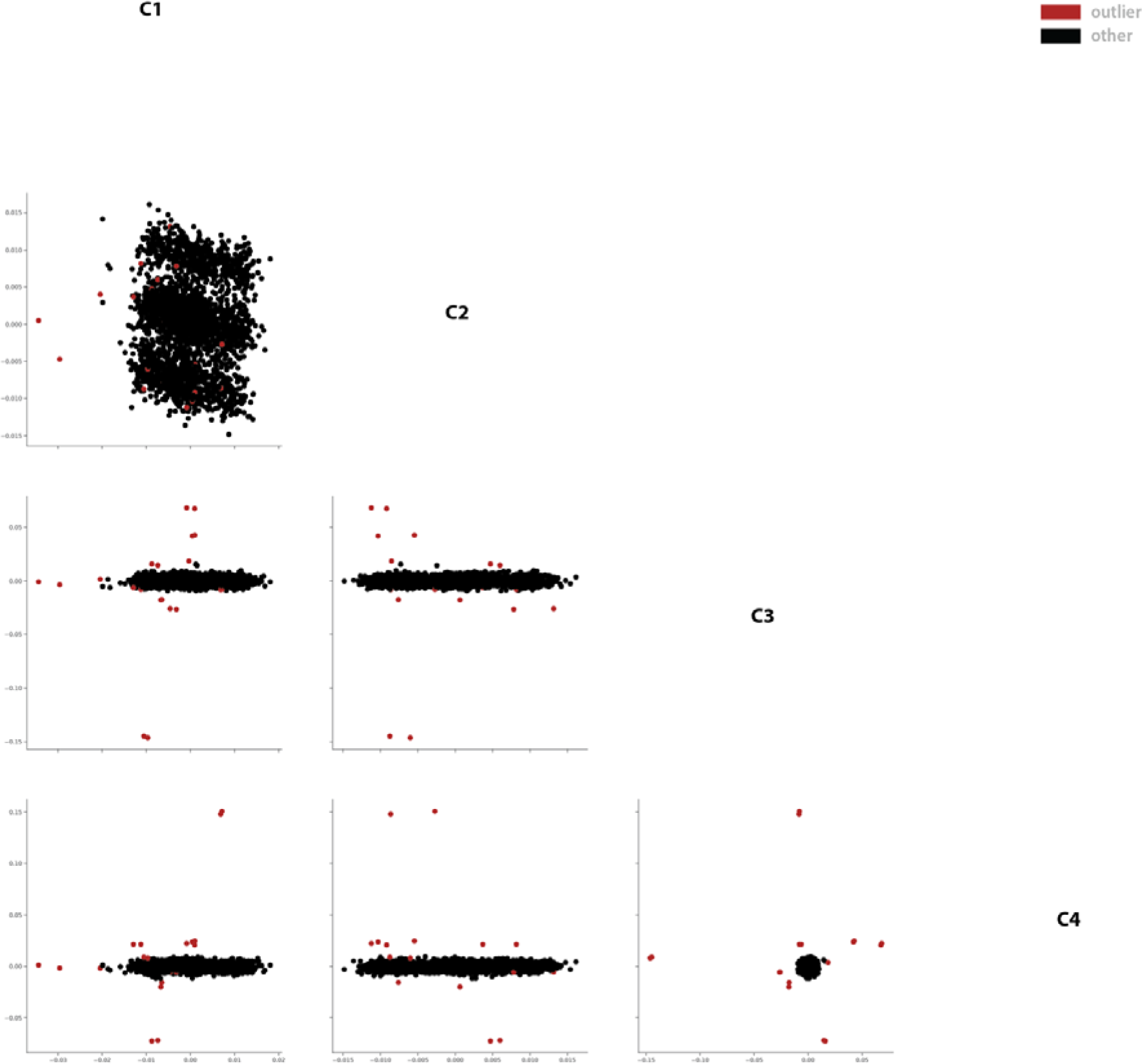
Blood genotype MDS outlier removal. *Exclusion of genotype outlier samples in blood. Samples that have absolute z-score >3 for any of the first four genotype MDS components are excluded (colored in red)*.

**Fig. S15.**
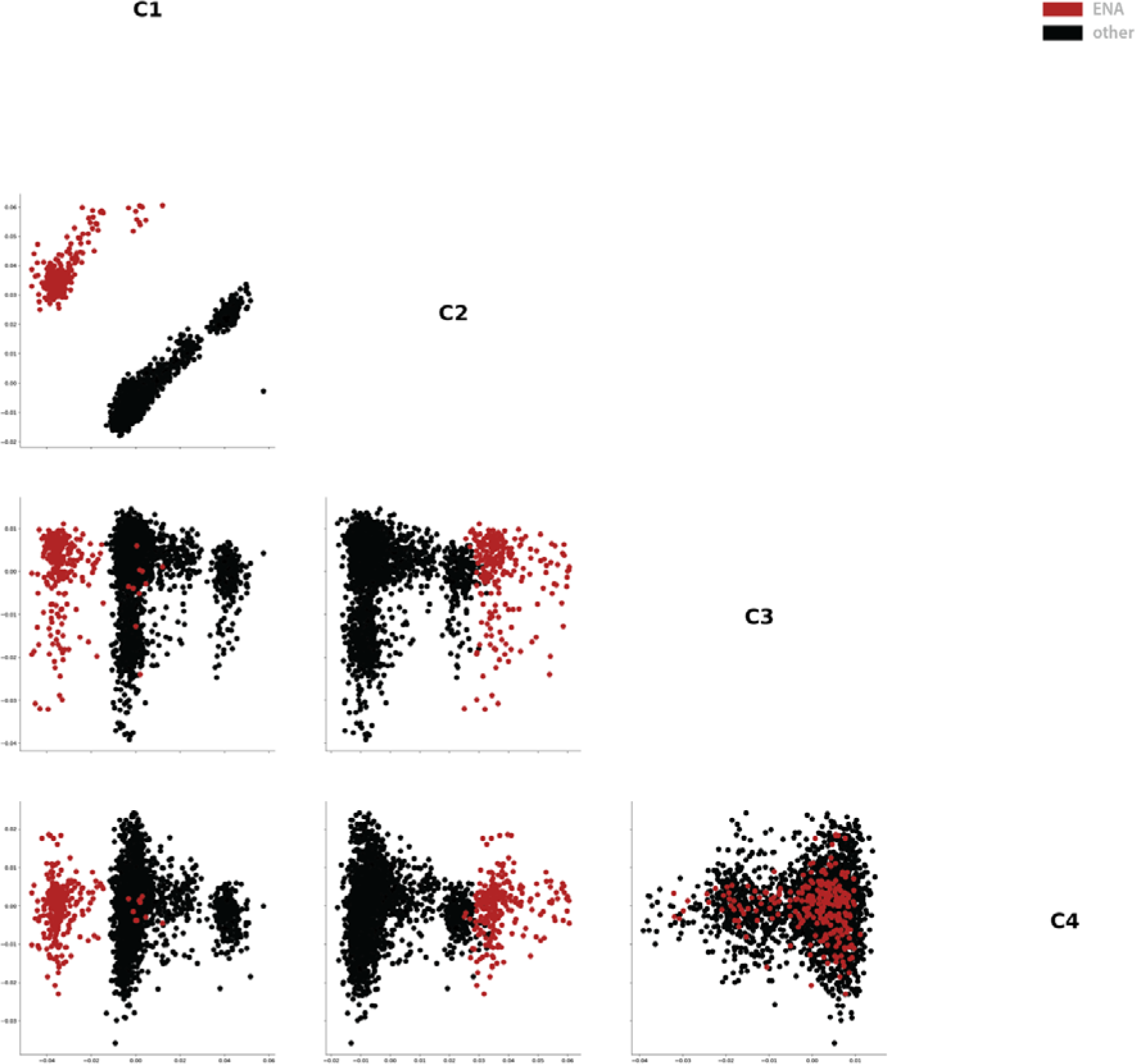
Brain genotype MDS outlier removal. *Exclusion of genotype outlier samples in brain. The ENA samples show to be a clear outlier (colored in red), probably because these genotypes were derived from sequencing data*.

**Fig. S16.**
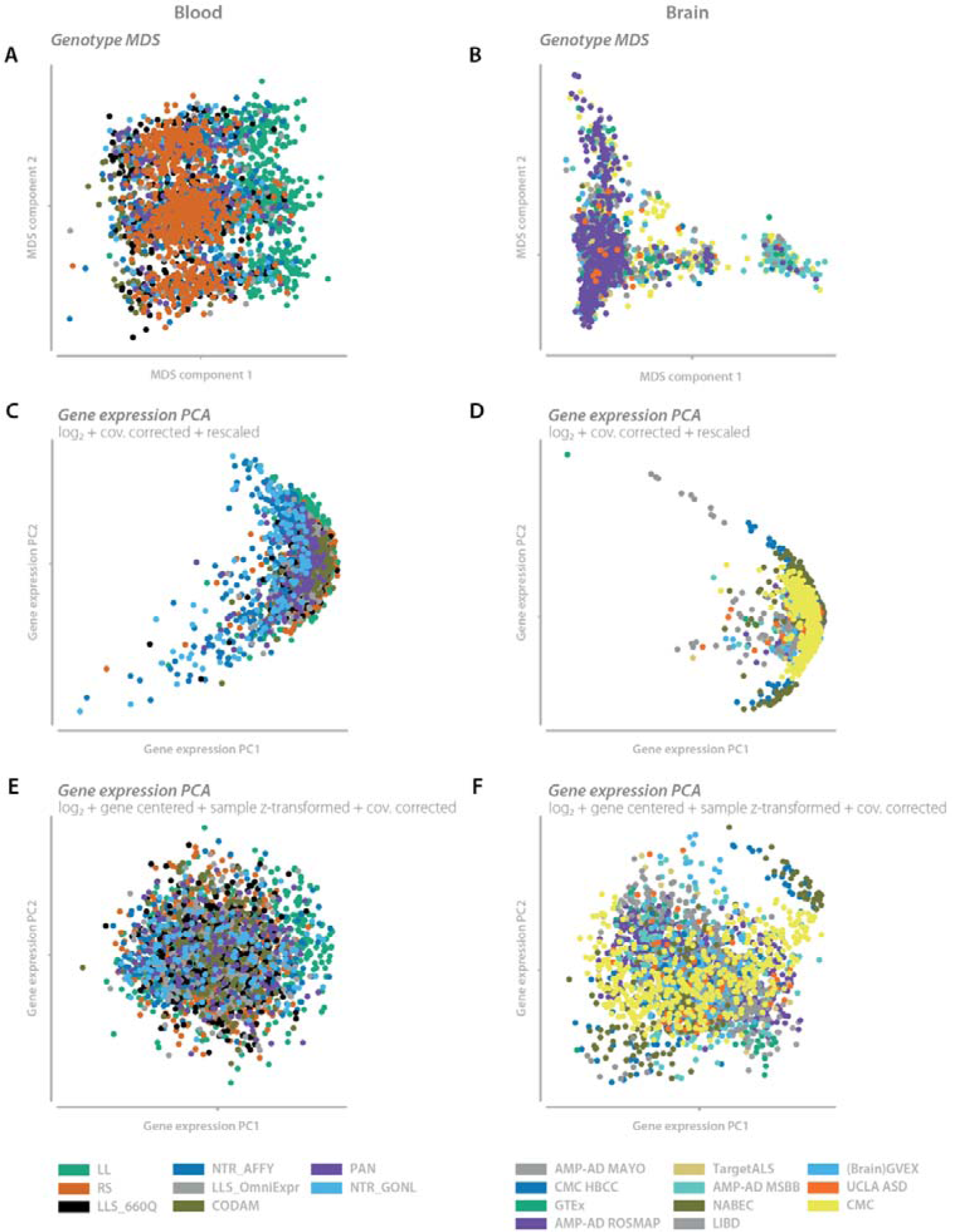
Genotype MDS and gene expression PCA for brain and blood. *Genotype and gene expression dimensional reduction plots for blood and brain after removal of outliers. (A, B) The first two genotype MDS components for blood (A) and brain (B). (C, D) The first two gene expression PCA components on TMM normalized, log_2_ transformed, OLS covariates removed (sex, four genotype MDS, and dataset indicator variables), and log_2_ mean and standard deviation returned data for blood (C) and brain (D). (E, F) The first two gene expression PCA components on TMM normalized, log_2_ transformed, gene centered, sample z-score transformed, OLS covariates removed (sex, four genotype MDS, and dataset indicator variables) data for blood (E) and brain (F)*.

Table S1. Number of interaction eQTL per expression PC or PIC in blood and brain Number of interaction-eQTL (ieQTLs) per expression PC or PICS in blood and brain. **Sheet blood**: contains the results for the analysis in blood. **Sheet brain**: contains the results for the analysis in brain. **Columns starting with PIC**: number of ieQTLs with PICs. **Columns starting with PC**: number of ieQTLs with expression PCs. **Columns ending with default**: interaction analysis where each component is tested irrespective of the others. **Columns ending with conditional**: interaction analysis where each previous component and its interaction with genotype have been removed from the expression data prior to mapping ieQTLs.

Table S2. Blood conditional PIC ieQTL summary statistics *Interaction eQTL summary statistics for blood using conditional analysis (i.e.: PIC1 is removed before evaluating PIC2 etc.). Sheet ‘SNPAndGeneAnnotation’ contains the eQTL annotation information. **SNP**: eQTL SNP. **Alleles**: SNP alleles. **affect allele**: the allele to which the betas are directed. **MAF**: the minor allele frequency **HW p-value**: the Hardy-Weinberg equilibrium p-value. **gene**: the ENSEMBL gene id. **symbol**: the HGNC gene symbol. **N**: the sample size. Each subsequent sheet contains interaction summary statistics per PIC. **SNP**: eQTL SNP. **gene**: the ENSEMBL gene id. **beta-intertaction**: the linear regression betas for the interaction term. **Std- interaction**:: the linear regression standard error for the interaction term. **p-value**: the p-value for the interaction term. **BH-FDR**: the Benjamini-Hochberg FDR for the interaction term*.

Table S3. Brain conditional PIC ieQTL summary statistics *Interaction eQTL summary statistics for brain using conditional analysis (i.e.: PIC1 is removed before evaluating PIC2 etc.). Sheet ‘SNPAndGeneAnnotation’ contains the eQTL annotation information.. **SNP**: eQTL SNP. **Alleles**: SNP alleles. **affect allele**: the allele to which the betas are directed. **MAF**: the minor allele frequency **HW p-value**: the Hardy-Weinberg equilibrium p-value. **gene**: the ENSEMBL gene id. **symbol**: the HGNC gene symbol. **N**: the sample size. Each subsequent sheet contains interaction summary statistics per PIC. **SNP**: eQTL SNP. **gene**: the ENSEMBL gene id.**beta-intertaction**: the linear regression betas for the interaction term. **Std- interaction**: the linear regression standard error for the interaction term. **p-value**: the p-value for the interaction term. **BH-FDR**: the Benjamini-Hochberg FDR for the interaction term*.

Table S4. Blood and brain gene expression vs PIC correlations *Pearson correlation summary statistics between gene expression and PICs for blood and brain. For gene expression we used TMM log_2_ transformed, gene cantered, sample z-score transformed, OLS covariates removed (sex, four genotype MDS, and dataset indicator variables), and finally forced the gene expression levels into a normal distribution per dataset by ranking with ties. Sheet ‘Blood’ contains the blood data, sheet ‘Brain’ contains the brain data. **gene**: the ENSEMBL gene id. **symbol**: the HGNC gene symbol. **Average TMM log2 expression**: the average TMM log_2_ expression of the gene for the included samples. **Columns ending with r**: the Pearson correlation coefficient rounded on six significant digits*.

Table S5. ToppGene gene list enrichment *ToppGene gene set enrichment for blood (sheets starting with blood) and brain (sheets starting with brain), for the ToppCell and Pathway databases, and performed for the top genes correlating genes (maximum of 200; positive / negative separated) as well as the eQTL genes that interact with a given PIC (sheets ending with ieQTL genes). **PIC**: the PIC. **Name**: the name of the ToppCell record or pathway. **p-value**: the p-value for the enrichment. **BH-FDR**: the Benjamini-Hochberg corrected p-value for the enrichment. **Number of genes in category**: the number of genes included in either ToppCell or Pathway annotations. **Number of input genes in category**: the number of PIC correlating or interacting genes that overlap with the genes in ToppCell or Pathway. **Number of genes in annotation**: the number of genes included in the individual ToppCell record or pathway*. ***Number of genes from input****: the number of PIC correlating or interacting genes that overlap with the individual ToppCell record or pathway. **HGNC symbols of genes from input**: the gene symbols for the PIC correlating or interacting genes. **Source**: the source of the ToppCell or Pathway annotation. **URL**: the URL of the ToppCell or Pathway annotation. **Correlation direction**: if the Pearson correlation coefficient between the PIC and the gene expression is below (negative) or above (positive) zero*.

Table S6. Replication of the MetaBrain EUR ieQTLs in AFR samples. *Sheet Summary: the replication statistics per PIC. **N**: number of eQTLs with a significant interaction in EUR. **Pearson r**: Pearson correlation coefficient between interaction t-values. **Concordance**: percentage of ieQTLs with the same direction of effect. **Rb**: Rb statistic over the interaction beta. **Pi1**: π1 statistic over the interaction p-values. Sheet Data: **Gene**: eQTL gene ENSEMBL ID. **Gene symbol**: eQTL gene symbol. **SNP**: eQTL SNP. **Alleles**: SNP alleles. **Allele assessed**: the allele to which the betas are directed. **Columns ending with N**: the sample size. **Columns ending with HW pval**: the Hardy-Weinberg equilibrium p-value. **Columns ending with Minor allele**: the minor allele. **Columns ending with MAF**: the minor allele frequency. **Columns ending with tvalue**: beta divided by the standard deviation for the genotype x PIC interaction term. **Columns ending with pvalue**: p-value for the ieQTL. **Columns ending with FDR**: the Benjamini-Hochberg FDR. For the replication datasets this FDR is calculated over the discovery significant (BH-FDR*≤*0.05) rows for each PIC*.

