## Supplementary Note for "Unbiased identification of unknown cellular and environmental factors that mediate eQTLs using principal interaction component analysis"

**PICS identified with PICALO are robust with respect to the initial guess used for optimization**

Since EM algorithms can yield results that depend on the choice of the initial starting position, we evaluated if the outcome of PICALO was dependent on the initial guess used for optimization (by default the initial guess with most significant ieQTLs). For each of the first five PICs in blood, we reperformed the analysis while using each of the first 25 expression PCs as the initial guess for optimization and compared the outcomes. For PIC1 we found that irrespective of the initial guess, all 25 PICALO analyses resulted in near-identical outcomes (average pair-wise Pearson r=0.9997; Fig. S5A). We note that this was independent of the number of significant ieQTLs that was used to start the optimization (between 4 and 2,670, depending on the PC; Fig. S5B). We then adjusted the expression data for PIC1 and its interaction effect with eQTLs and evaluated PIC2. Similarly, we observed a robust outcome of PICALO for 21 out of the 25 expression PCs used as initial guess, with an average pair-wise Pearson r=0.9946. The remaining four PCs resulted in two distinct groups of PICALO outcomes: the first group (PC13 and PC20) had a Pearson r of 0.65 and £0.05 with the other outcomes, while the second group (PC14 and PC21) had a Pearson r of 0.44 and £0.01 with others. Each of these groups had a smaller number of significant ieQTLs than the original PIC2 (Number of ieQTLs; PIC2 equivalents: 1,053 ±27, first group: 209 ±10, second group: 22 ±8). This observation confirms that, like other EM methods, PICALO may not always return the global optimum but may instead converge to a local optimum. Like for PIC1, we found no relationship between the number of eQTLs interacting with the starting PC and the resulting PIC (Fig. S5B). Since this step yielded multiple solutions, we selected the PIC with the largest number of ieQTLs before optimization as PIC2, identical to a normal PICALO procedure. For the subsequent PICs we observed that an increasing number of uncorrelated PICALO outcomes were identified dependent on the initial guess that was used for optimization. Notably, we observed that the four local optimums identified in the PIC2 analyses are rediscovered as local optimums in PIC3, an example of which is shown in Fig. S5C. In other words: the influence of the initial guess on the resulting PIC increases as the local optimums in the data capture decreasing proportions of interaction variance. These observations suggest that PICs capture unique effects that can be robustly identified using PICALO, but that the order in which PICs are identified can be dependent on the initial guess.

**Supplementary Note 2**

**BIOS Consortium (Biobank-based Integrative Omics Study) – Author information**

**Management Team**
Bastiaan T. Heijmans (chair)^1^, Peter A.C. ’t Hoen^2^, Joyce van Meurs^3^, Rick Jansen^5^, Lude Franke^6^.

**Cohort collection**
Dorret I. Boomsma^7^, René Pool^7^, Jenny van Dongen^7^, Jouke J. Hottenga^7^ (Netherlands Twin Register); Marleen MJ van Greevenbroek^8^, Coen D.A. Stehouwer^8^, Carla J.H. van der Kallen^8^, Casper G. Schalkwijk^8^ (Cohort study on Diabetes and Atherosclerosis Maastricht); Cisca Wijmenga^6^, Lude Franke^6^, Sasha Zhernakova^6^, Ettje F. Tigchelaar^6^ (LifeLines Deep); P. Eline Slagboom^1^, Marian Beekman^1^, Joris Deelen^1^, Diana van Heemst^9^ (Leiden Longevity Study); Jan H. Veldink^10^, Leonard H. van den Berg^10^ (Prospective ALS Study Netherlands); Cornelia M. van Duijn^4^, Bert A. Hofman^11^, Aaron Isaacs^4^, André G. Uitterlinden^3^ (Rotterdam Study).

**Data Generation**
Joyce van Meurs (Chair)^3^, P. Mila Jhamai^3^, Michael Verbiest^3^, H. Eka D. Suchiman^1^, Marijn Verkerk^3^, Ruud van der Breggen^1^, Jeroen van Rooij^3^, Nico Lakenberg^1^.

**Data management and computational infrastructure**
Hailiang Mei (Chair)^12^, Maarten van Iterson^1^, Michiel van Galen^2^, Jan Bot^13^, Dasha V. Zhernakova^6^, Rick Jansen^5^, Peter van ’t Hof^12^, Patrick Deelen^6^, Irene Nooren^13^, Peter A.C. ’t Hoen^2^, Bastiaan T. Heijmans^1^, Matthijs Moed^1^.

**Data Analysis Group**
Lude Franke (Co-Chair)^6^, Martijn Vermaat^2^, Dasha V. Zhernakova^6^, René Luijk^1^, Marc Jan Bonder^6^, Maarten van Iterson^1^, Patrick Deelen^6^, Freerk van Dijk^14^, Michiel van Galen^2^, Wibowo Arindrarto^12^, Szymon M. Kielbasa^15^, Morris A. Swertz^14^, Erik. W van Zwet^15,^ Rick Jansen^5^, Peter-Bram ’t Hoen (Co-Chair)^2^, Bastiaan T. Heijmans (Co-Chair)^1^.

1. Molecular Epidemiology, Department of Biomedical Data Sciences, Leiden University Medical Center, Leiden, The Netherlands
2. Department of Human Genetics, Leiden University Medical Center, Leiden, The Netherlands
3. Department of Internal Medicine, ErasmusMC, Rotterdam, The Netherlands
4. Department of Genetic Epidemiology, ErasmusMC, Rotterdam, The Netherlands
5. Department of Psychiatry, VU University Medical Center, Neuroscience Campus Amsterdam, Amsterdam, The Netherlands
6. Department of Genetics, University of Groningen, University Medical Centre Groningen, Groningen, The Netherlands
7. Department of Biological Psychology, VU University Amsterdam, Neuroscience Campus Amsterdam, Amsterdam, The Netherlands
8. Department of Internal Medicine and School for Cardiovascular Diseases (CARIM), Maastricht University Medical Center, Maastricht, The Netherlands
9. Department of Gerontology and Geriatrics, Leiden University Medical Center, Leiden, The Netherlands
10. Department of Neurology, Brain Center Rudolf Magnus, University Medical Center Utrecht, Utrecht, The Netherlands
11. Department of Epidemiology, ErasmusMC, Rotterdam, The Netherlands
12. Sequence Analysis Support Core, Department of Biomedical Data Sciences, Leiden University Medical Center, Leiden, The Netherlands
13. SURFsara, Amsterdam, the Netherlands
14. Genomics Coordination Center, University Medical Center Groningen, University of Groningen, Groningen, the Netherlands
15. Medical Statistics, Department of Biomedical Data Sciences, Leiden University Medical Center, Leiden, The Netherlands

**Software**

Python (v3.7.4)

R (v4.0.3)

**R packages**

qvalue^1^ (v2.15.0)

**Python packages**

numpy^2^ (v1.19.5), pandas^3^ (v1.2.1), scipy^4^ (v1.6.0), statsmodels^5^ (v0.12.2), matplotlib^6^ (v3.3.4), seaborn^7^ (v0.11.1), scikit-learn^8^ (v0.24.1), and upsetplot^9,10^ (v0.4.1)

10.	Nothman, J. UpSetPlot: Draw Lex et al.’s UpSet plots with Pandas and Matplotlib.
