## Supplementary figures and images for "Unbiased identification of unknown cellular and environmental factors that mediate eQTLs using principal interaction component analysis"

### Supplementary Figure 1

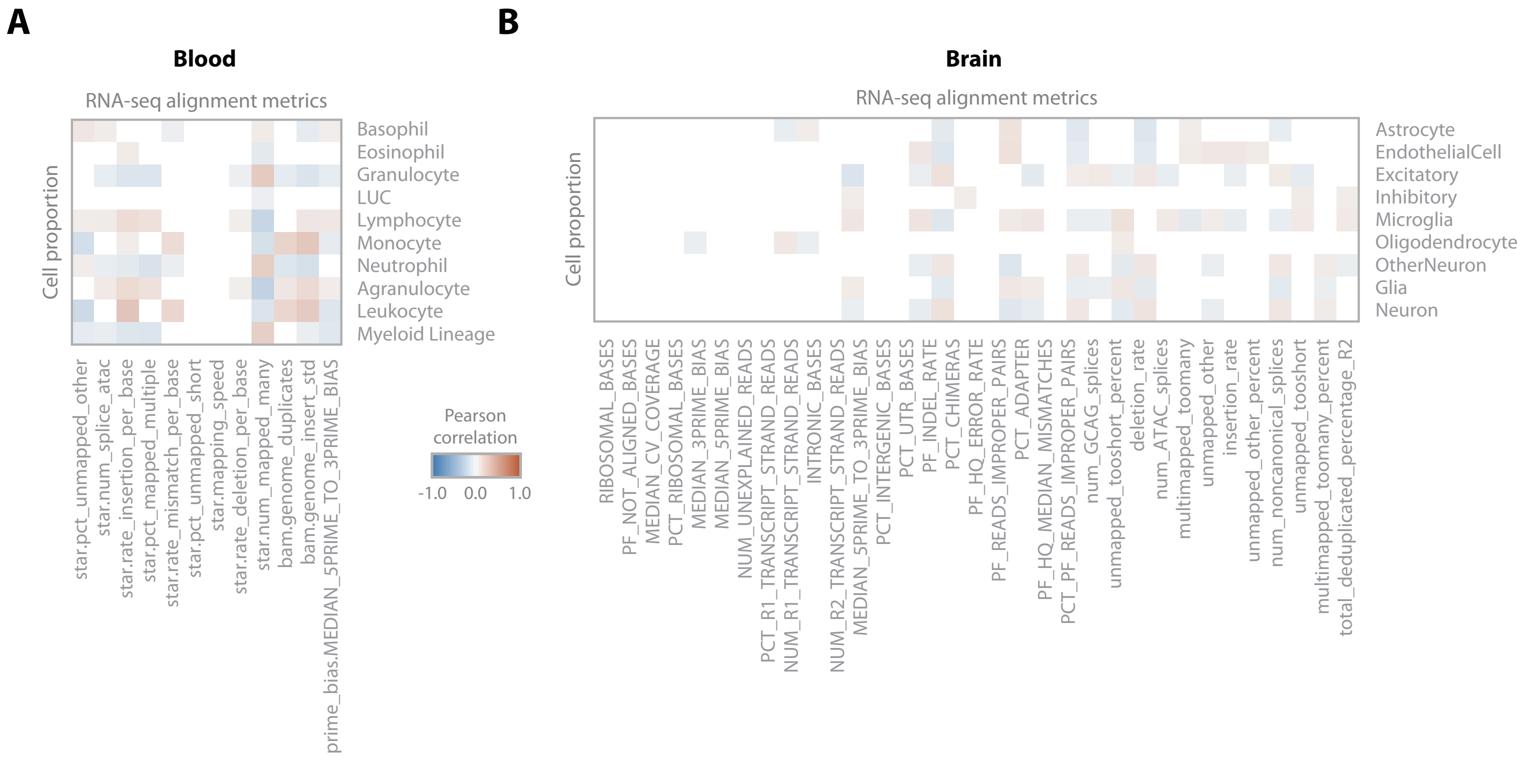

### Supplementary Figure 2

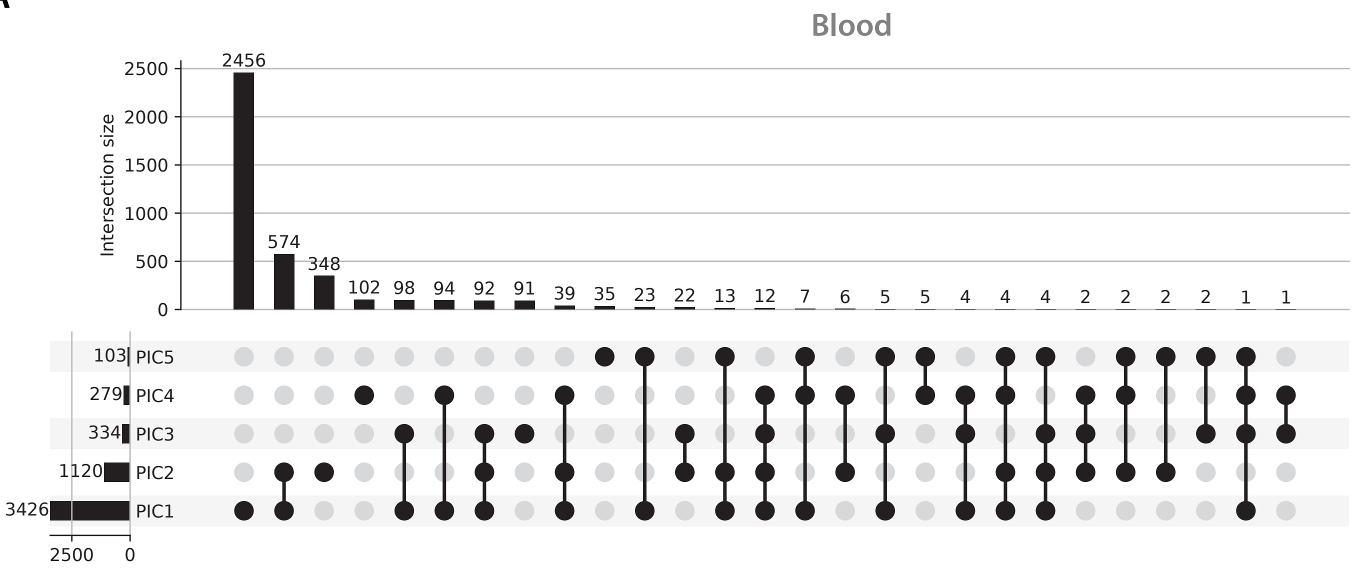

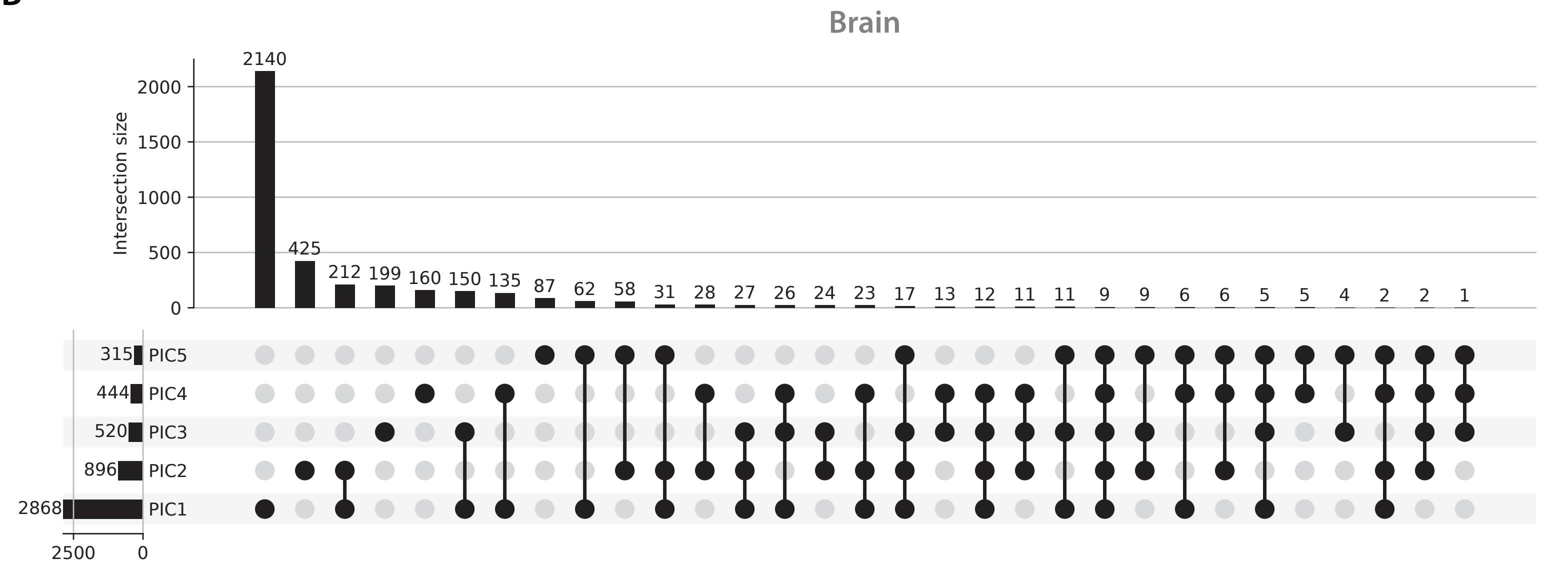

### Supplementary Figure 3

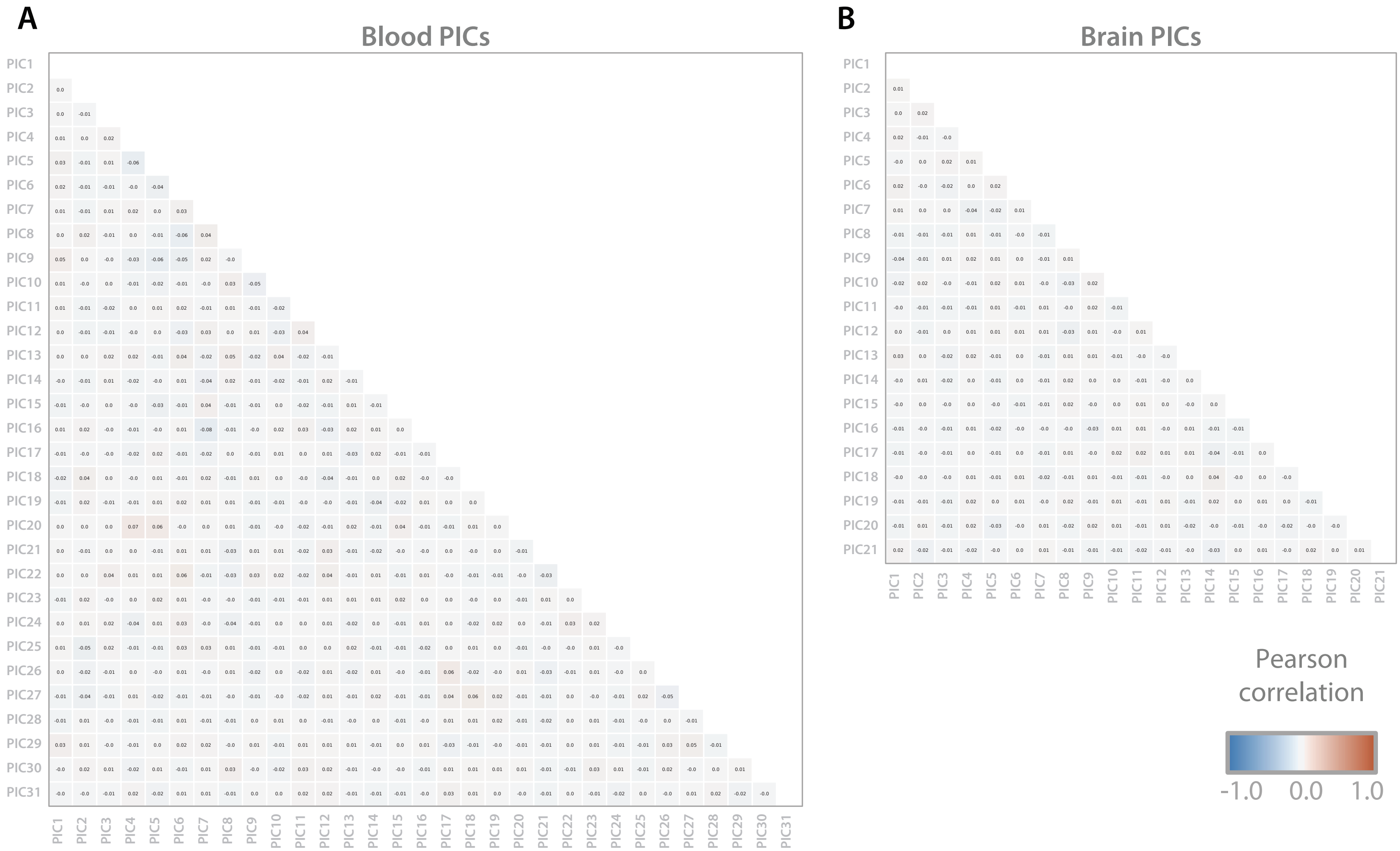

### Supplementary Figure 4

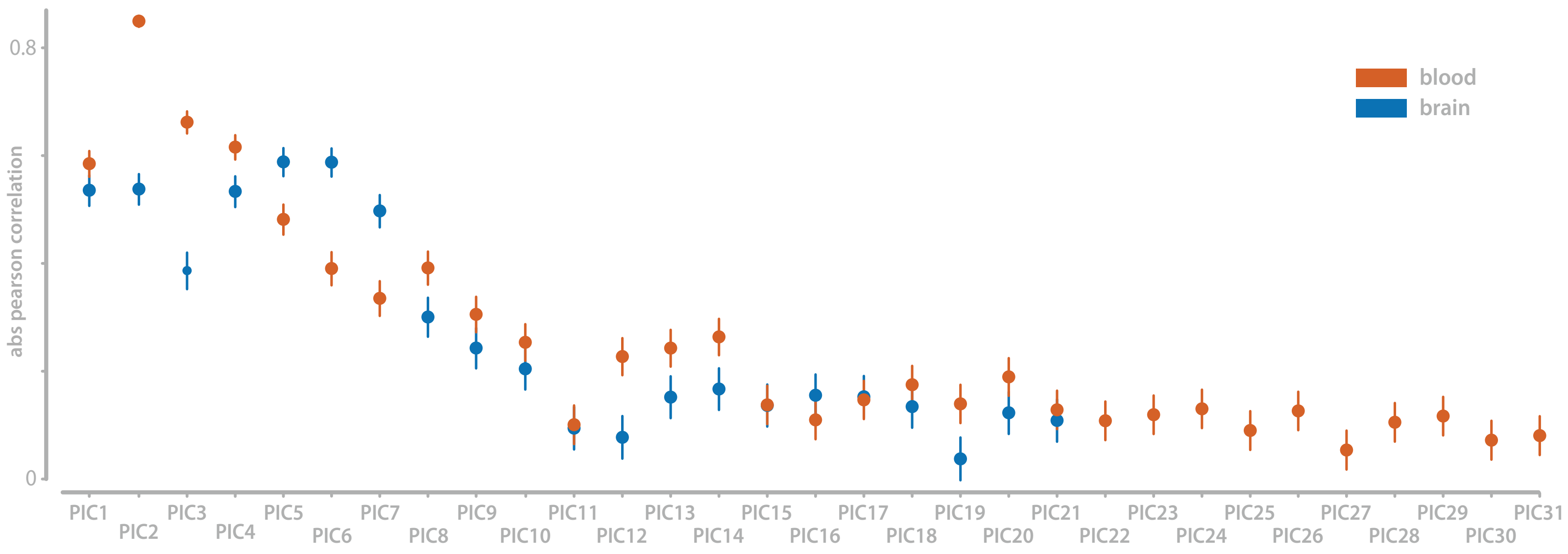

### Supplementary Figure 5

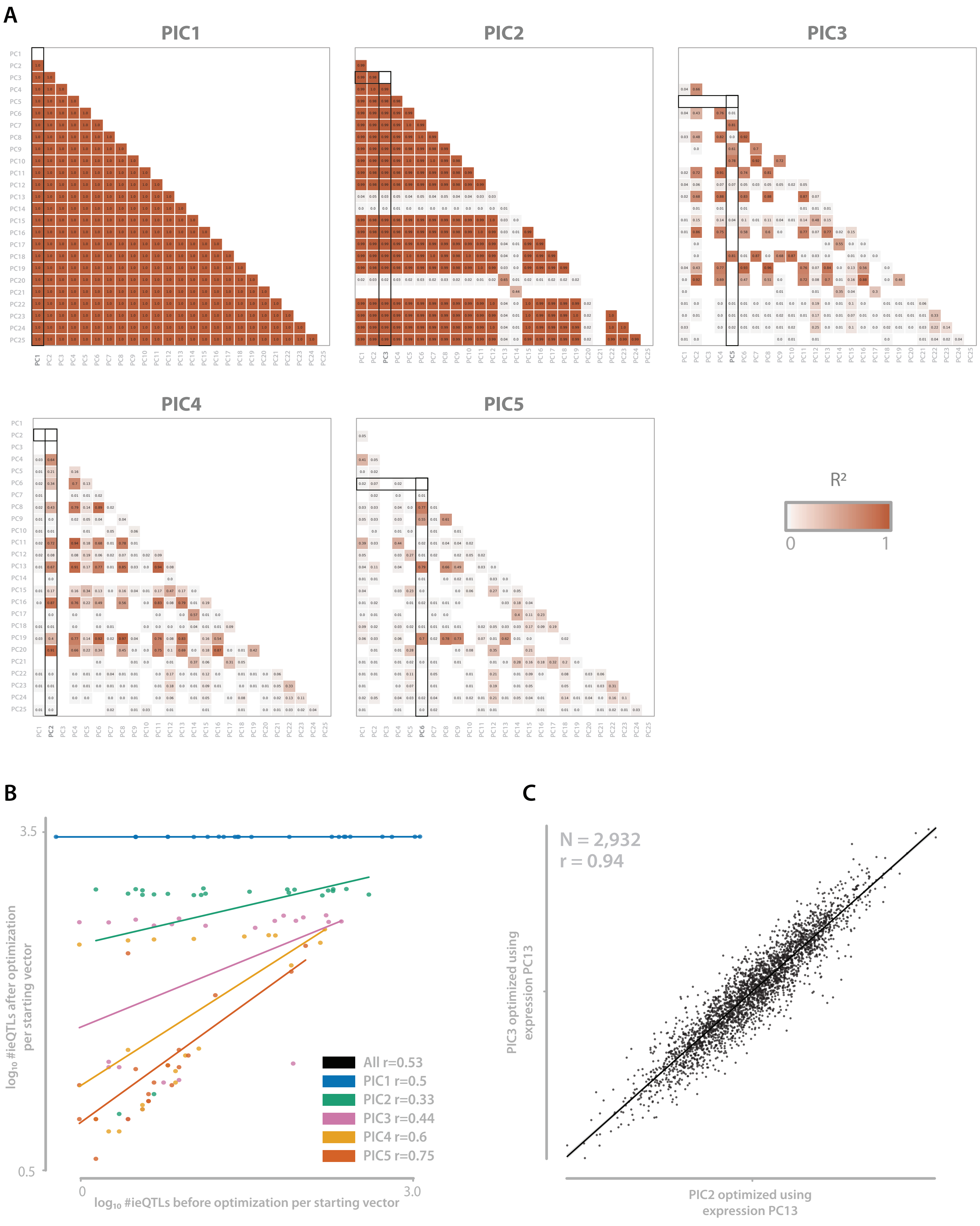

### Supplementary Figure 6

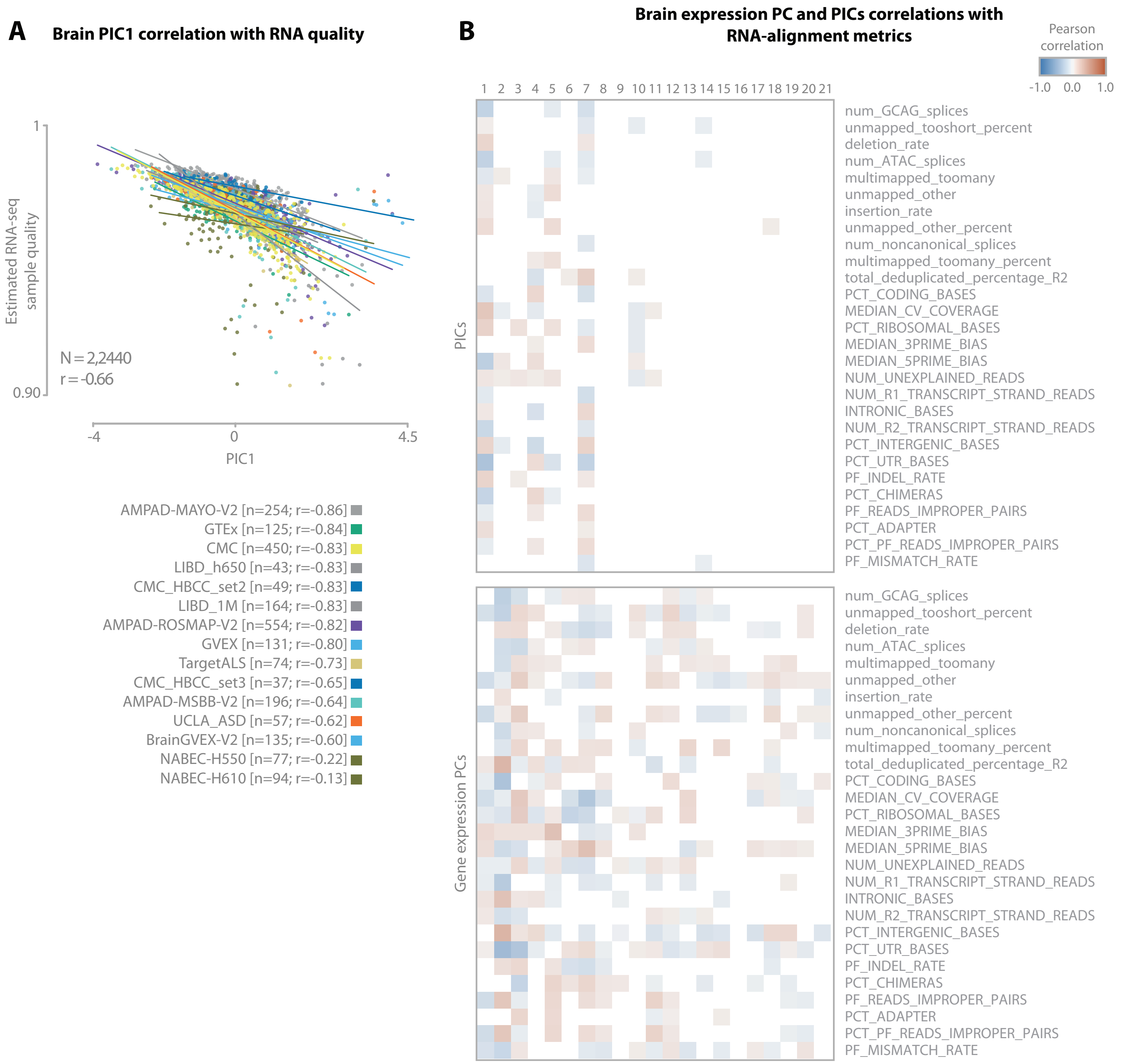

### Supplementary Figure 7

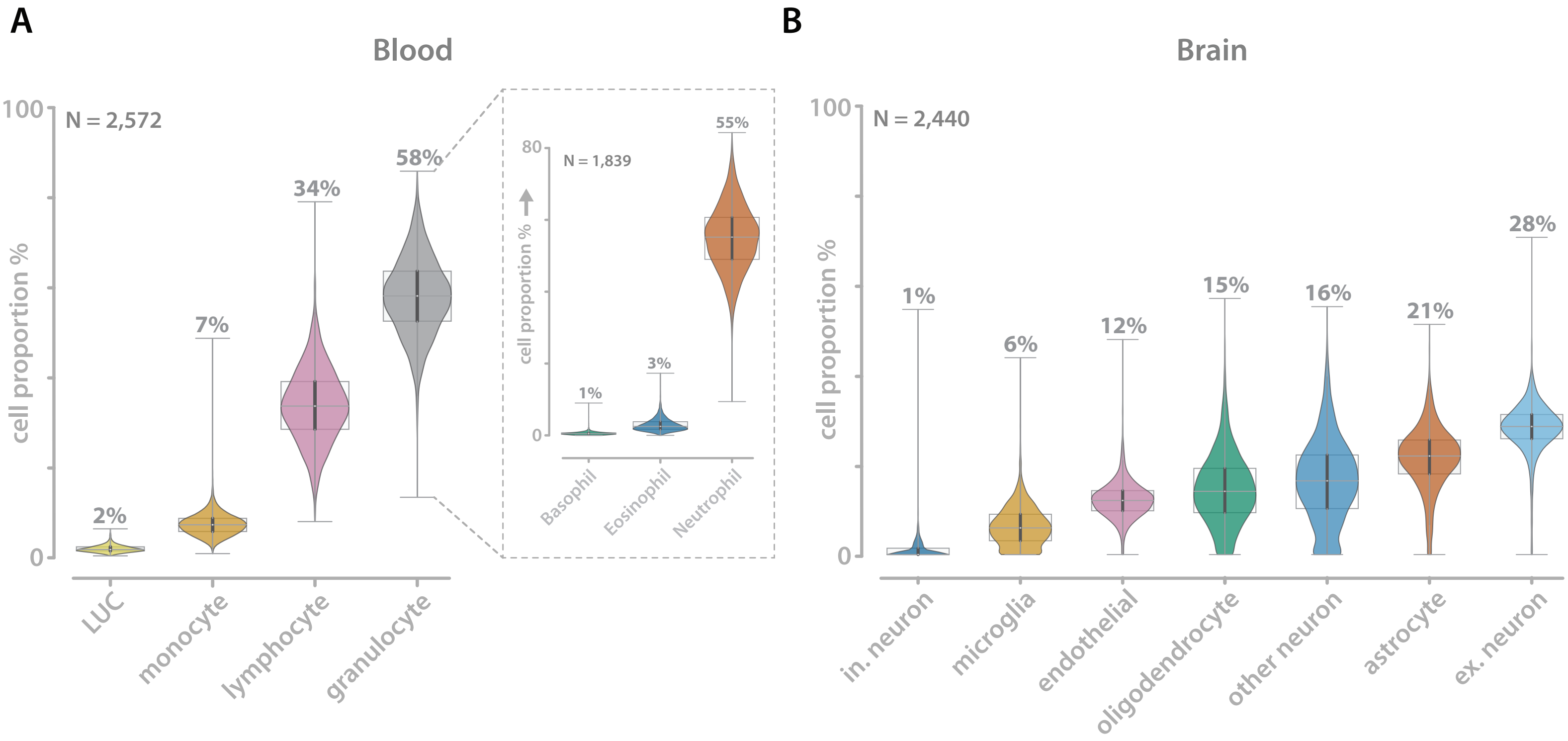

### Supplementary Figure 8

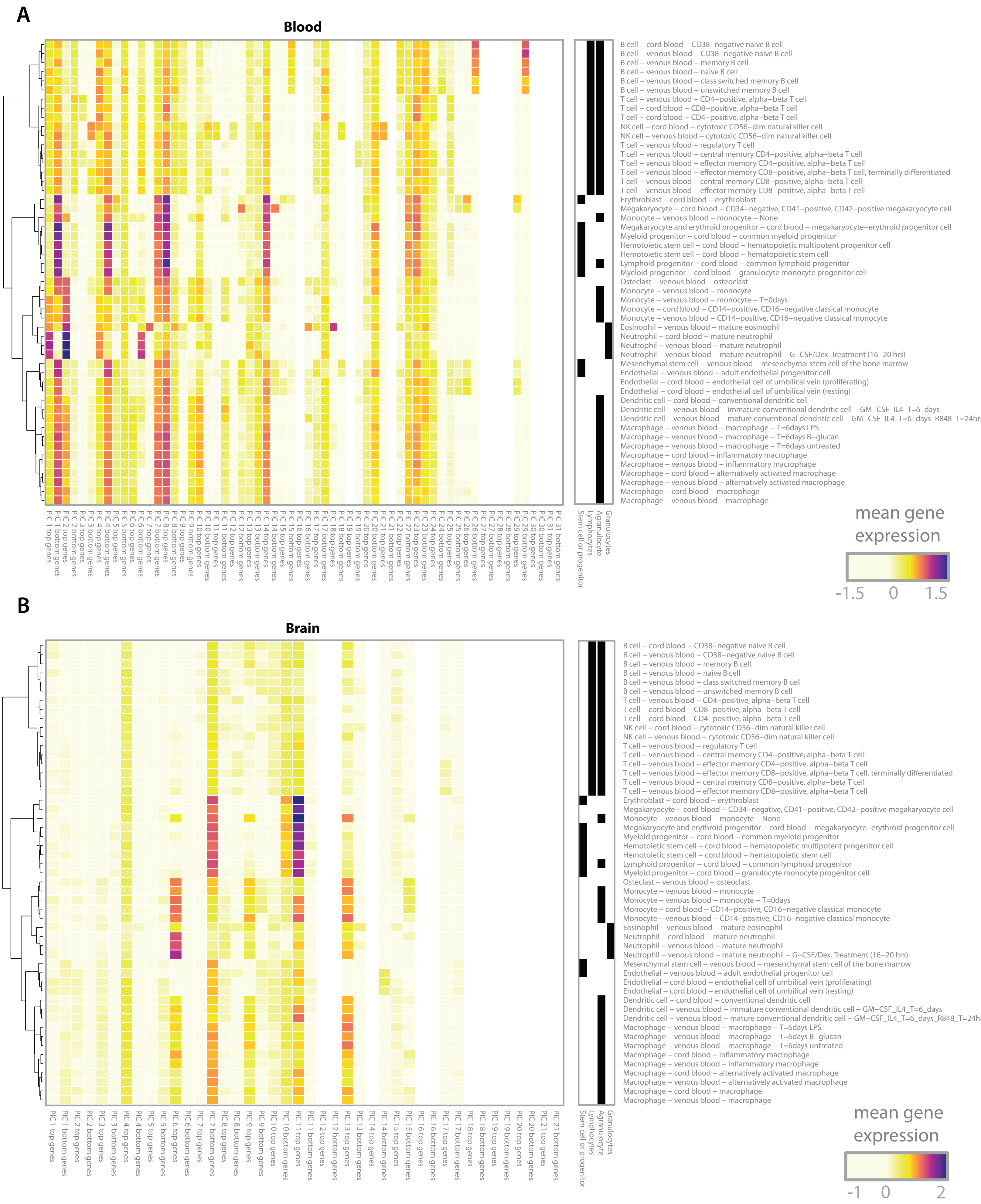

### Supplementary Figure 9

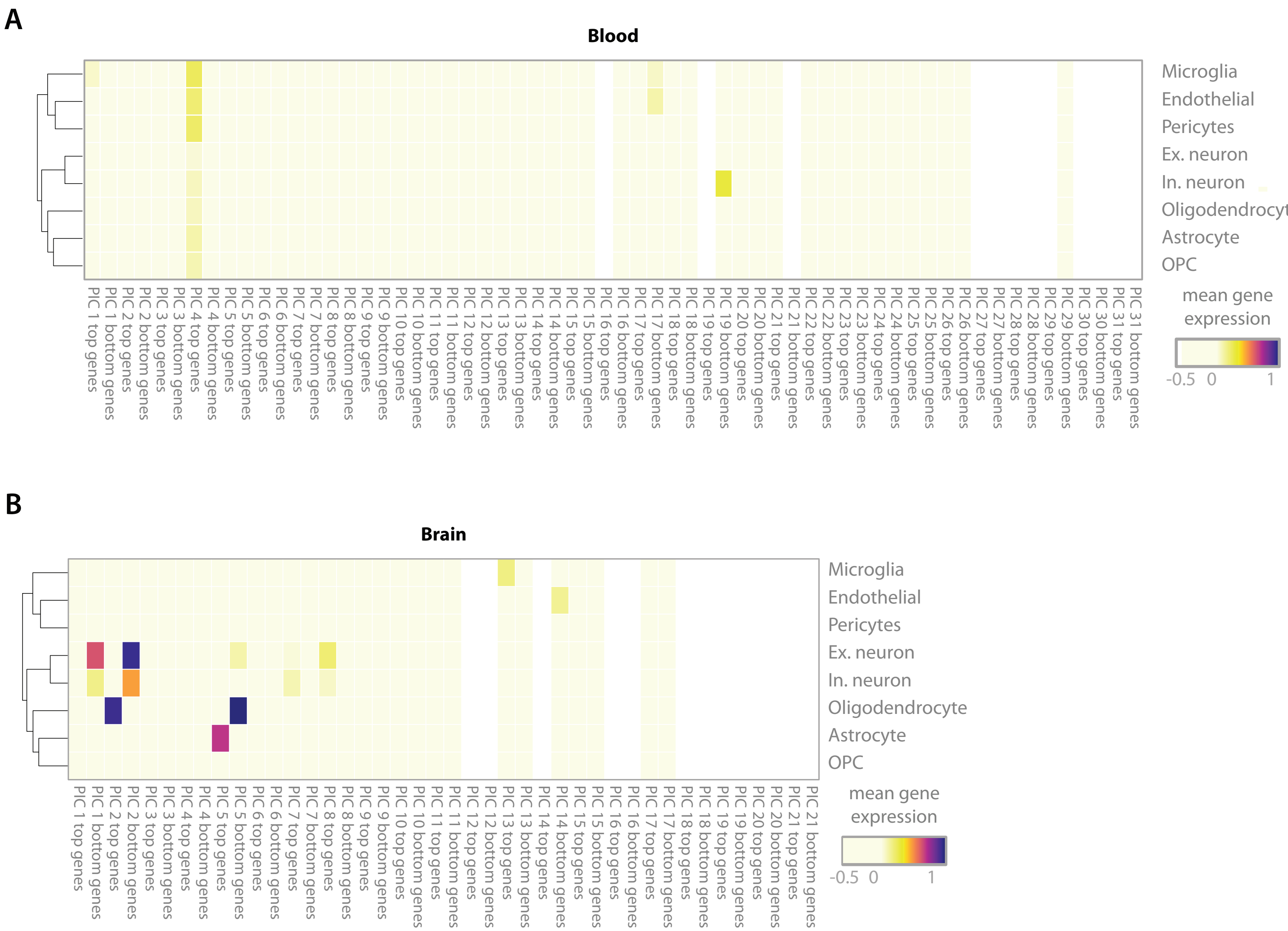

### Supplementary Figure 10

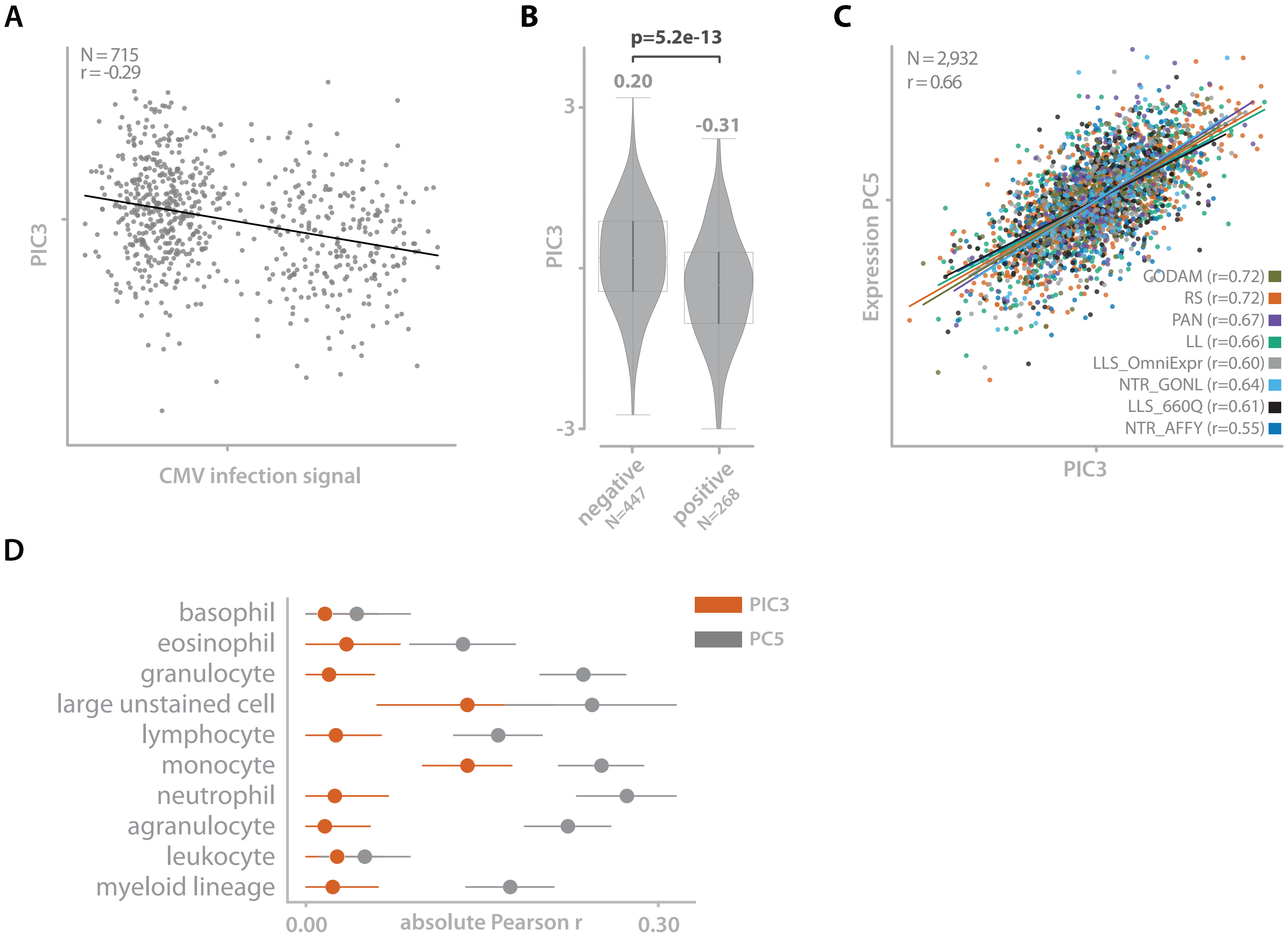

### Supplementary Figure 11

#### EUR PIC replication in AFR

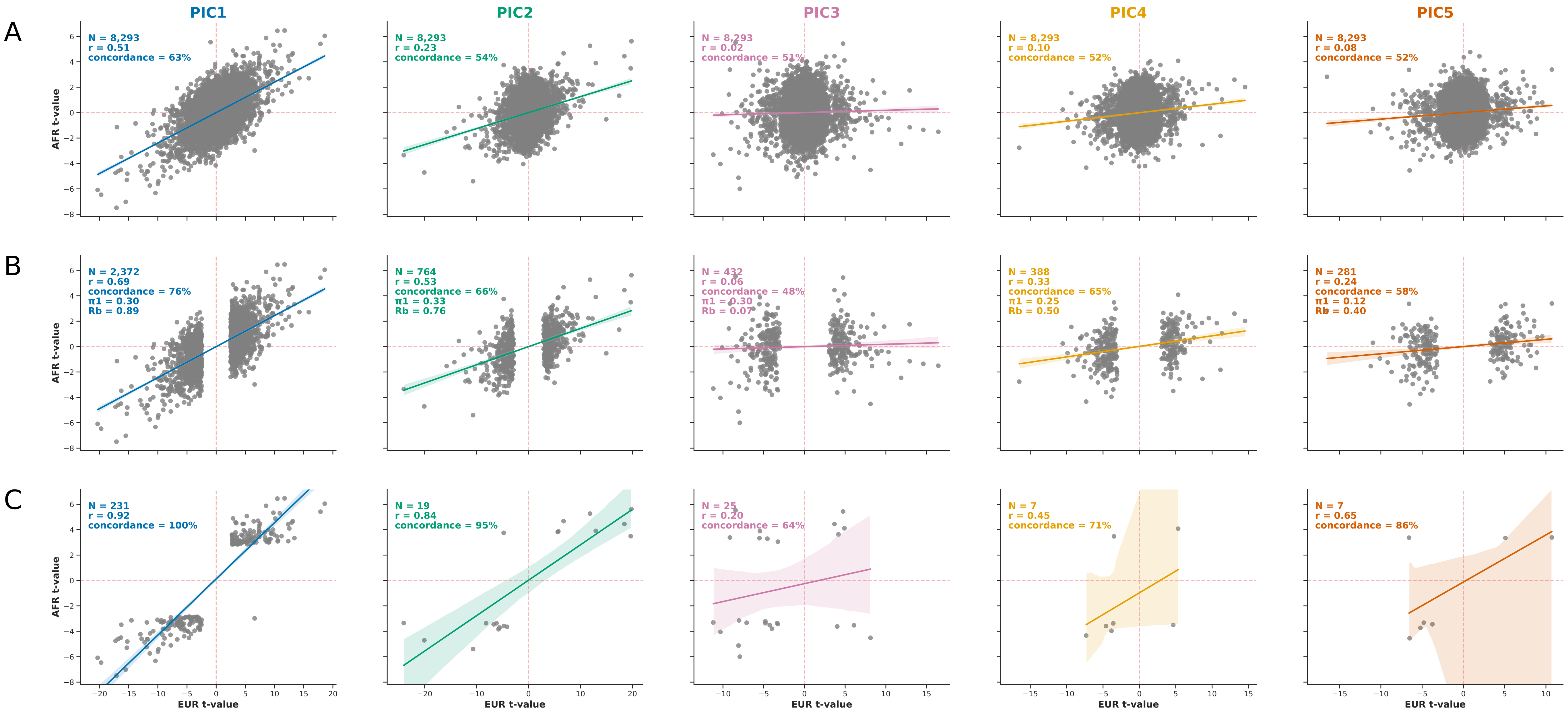

### Supplementary Figure 13

***Determine the optimal value for Jane Doe by finding the focus of the log likelihood function***

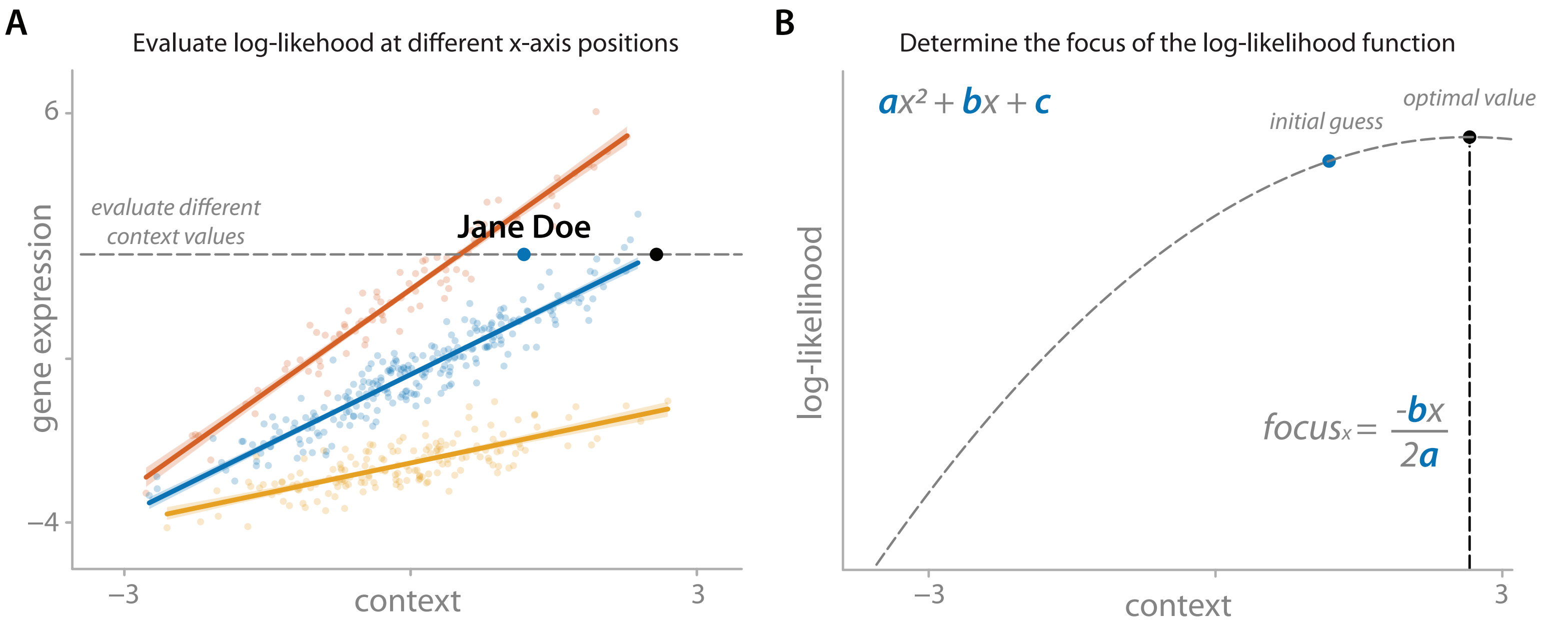

### Supplementary Figure 14

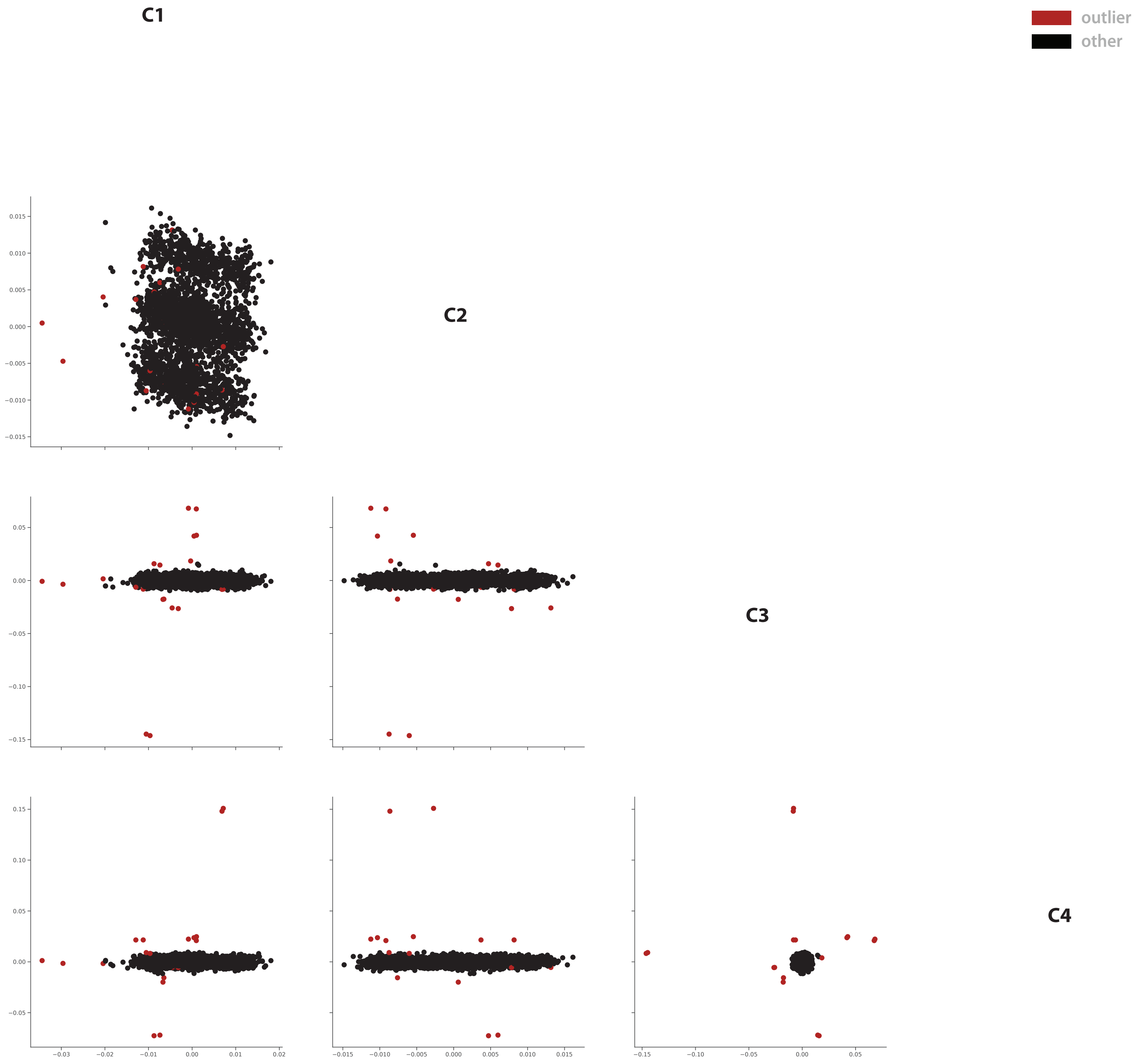

### Supplementary Figure 15

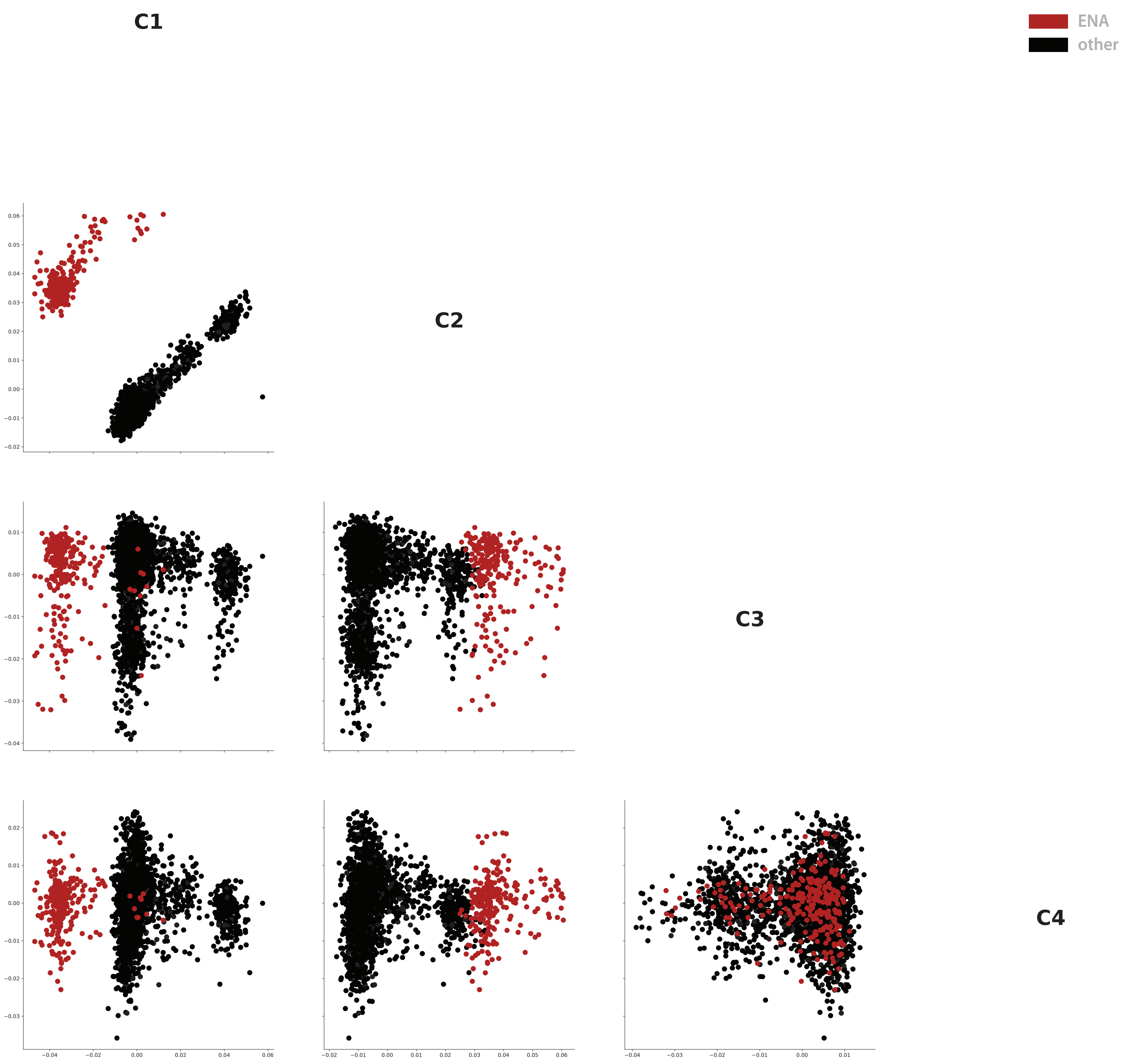

### Supplementary Figure 16

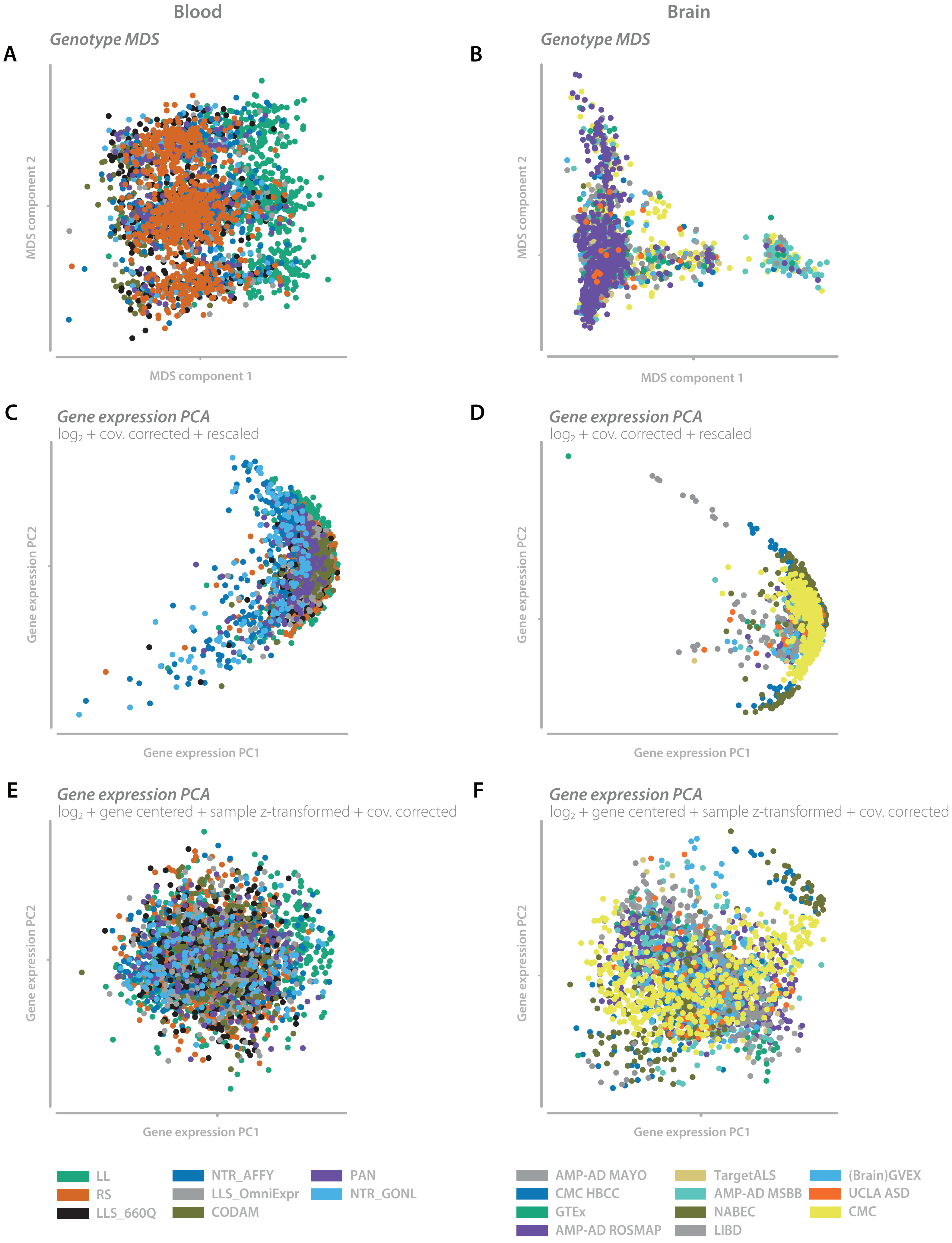
