## Supplementary Figure 12 for "Unbiased identification of unknown cellular and environmental factors that mediate eQTLs using principal interaction component analysis"

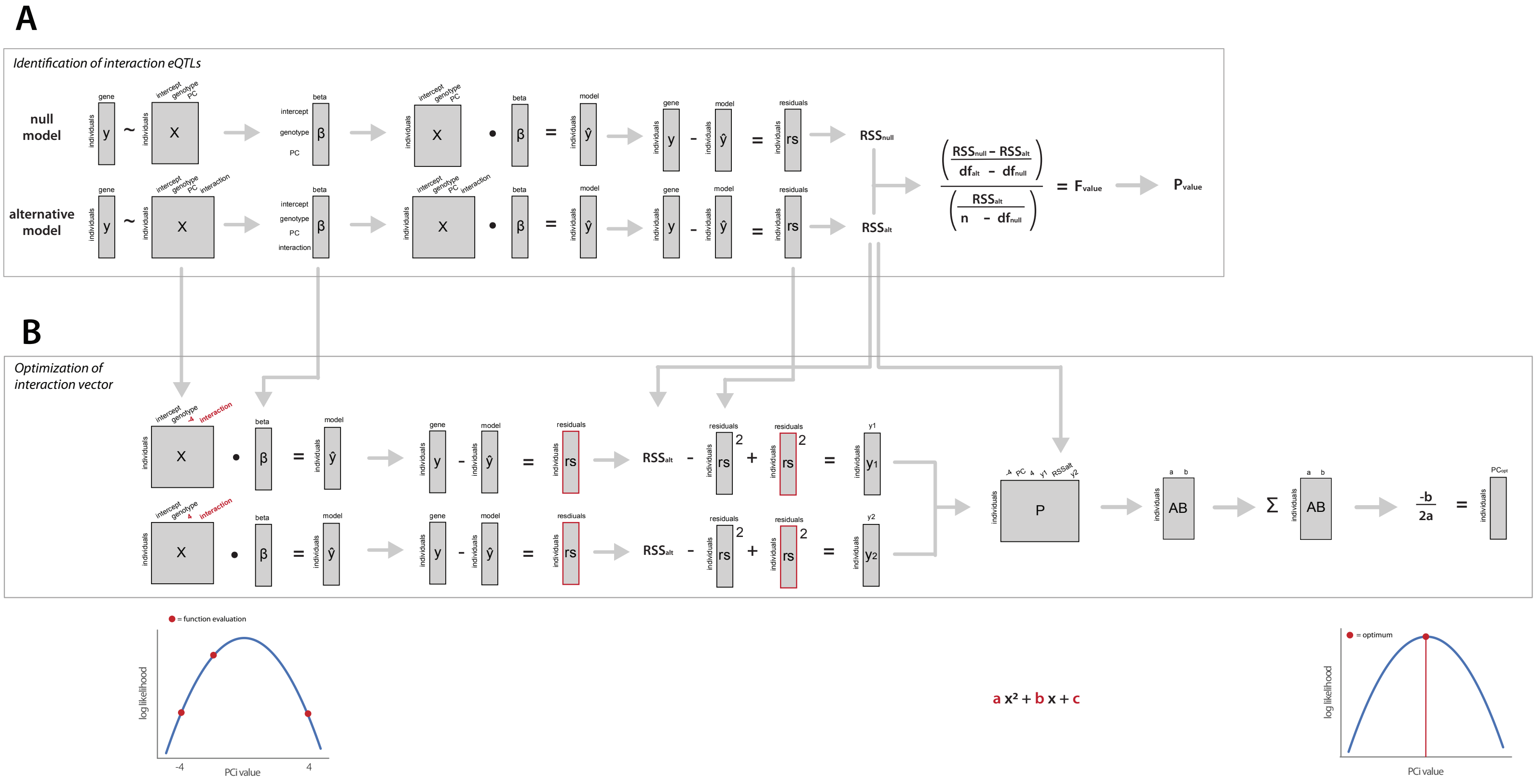


*Evaluate the log likelihood function on 3 unique points*

*Determine the coefcients of the likelihood parabola and join them to construct a joint function describing the join likelihood of a given context for all included interaction eQTLs*

*Determine the optimal value based on the focus x-axis position*
